# Stromal Interaction Molecule 1 Maintains β Cell Identity and Function in Female Mice through Preservation of G Protein-Coupled Estrogen Receptor 1 Signaling

**DOI:** 10.1101/2022.07.25.501429

**Authors:** Paul Sohn, Madeline R. McLaughlin, Preethi Krishnan, Chih-Chun Lee, Tatsuyoshi Kono, Carmella Evans-Molina

## Abstract

Loss of pancreatic β cell mass, identity, and function contribute to the development of diabetes. Here, we show that the endoplasmic reticulum (ER) calcium sensor, stromal interaction molecule 1 (STIM1), is critical for the maintenance of β cell function in female mice. When mice with β cell-specific deletion of STIM1 (STIM1Δβ) were challenged with high-fat diet, β cell dysfunction was observed in female, but not male, mice. Impaired glucose tolerance was accompanied by reductions in β cell mass, a concomitant increase in α cell mass, and significant reductions in the expression of markers of β cell maturity, including MafA and UCN3. Mechanistic assays demonstrated that the sexually dimorphic phenotype observed in STIM1Δβ mice was due in part to loss of signaling through the noncanonical 17-β estradiol receptor, GPER1. Together, these data suggest that STIM1 orchestrates pancreatic β cell function and identity through GPER1-mediated estradiol signaling.

## Introduction

Type 2 diabetes (T2D) affects over 537 million individuals worldwide and develops as a result of peripheral insulin resistance and pancreatic β cell dysfunction (Holman et al., 2008; Meier and Bonadonna, 2013). The vast majority of genetic variants associated with T2D risk are thought to act at the level of the β cell, and genetic studies indicate that certain “at-risk” individuals are more susceptible to β cell failure in settings of insulin resistance such as obesity and pregnancy (Florez et al., 2018; Herring and Oken, 2011). In addition to this genetic predisposition, systemic and local factors, including elevated levels of glucose, pro-inflammatory cytokines, and free fatty acids, directly contribute to impaired insulin secretion, β cell death, and dedifferentiation (Kahn et al., 2014; Moin and Butler, 2019).

Impairments in β cell calcium (Ca^2+^) signaling are linked to several molecular pathways that lead to β cell dysfunction. The fidelity of Ca^2+^ signaling requires the maintenance of steep concentration gradients, organized at both the cellular and organelle level. Extracellular Ca^2+^ concentrations are maintained in the range of 2.5 mM, while Ca^2+^ levels within the cytosol are kept at 100-250 nM under resting conditions. This low Ca^2+^ concentration is maintained by a series of molecular channels and pumps that translocate Ca^2+^ out of the cytosol and into the extracellular space or the endoplasmic reticulum (ER). In contrast to the low resting levels of Ca^2+^ observed within the cytosol, the Ca^2+^ concentration within the ER is estimated to be ~300-700 μM. Calcium within the ER lumen serves as an essential cofactor for molecular chaperones and foldases, including calnexin, calreticulin, and protein disulfide isomerase that ensure efficient protein synthesis and folding (Avezov et al., 2015; Wang et al., 2011).

In the pancreatic β cell, the ER serves as the dominant intracellular store of Ca^2+^ (Bygrave and Benedetti, 1996), and high levels of Ca^2+^ within the ER are maintained by the balance of ER Ca^2+^ uptake by the sarco-endoplasmic reticulum Ca^2+^ ATPase (SERCA) pump and ER Ca^2+^ release through the ryanodine receptors (RYR) and inositol 1,4,5-triphosphate receptors (IP3R) (Gilon et al., 2014; Santulli et al., 2017; Tong et al., 2016). In response to ER Ca^2+^ depletion, a process known as store-operated calcium entry (SOCE) replenishes ER Ca^2+^ levels through a family of channels referred to as store-operated or Ca^2+^ release-activated channels. During SOCE, decreased ER Ca^2+^ concentrations are detected by stromal interaction molecule 1 (STIM1), an ER Ca^2+^ sensor. STIM1 oligomerizes and translocates to the ER/plasmalemmal junctional regions (Roos et al., 2005), where it complexes with Orai Ca^2+^ channels to promote influx of Ca^2+^ from the extracellular space into the cytoplasm and ultimately into the ER lumen (Prakriya et al., 2006; Stathopulos et al., 2008).

SOCE has been accepted as a main pathway of Ca^2+^ entry into non-excitable cells, where it regulates a variety of functions including proliferation (Abdullaev et al., 2008), inflammation (Desvignes et al., 2015), lipid metabolism (Maus et al., 2017), cell senescence (Xu et al., 2015), and cell survival (Heise et al., 2010). The role of SOCE in excitable and secretory cells, such as pancreatic β cell, has been controversial; however, recent studies have implicated SOCE in glucose- and GPR40-mediated potentiation of insulin secretion (Usui et al., 2019). We have previously demonstrated reductions in STIM1 mRNA and protein expression in human islets from donors with T2D and in human islets and INS-1 β cells treated with pro-inflammatory cytokines and palmitate. Additionally, we have shown that STIM1 overexpression in human islets from donors with T2D improves insulin secretion (Kono et al., 2018). However, whether STIM1 loss impacts *in vivo* responses to metabolic stressors including diet-induced obesity remains unexplored.

In this study, we generated a transgenic mouse line with a pancreatic β cell-specific deletion of STIM1 (STIM1Δβ mice) and challenged these mice with a high-fat diet (HFD) to model diet-induced obesity. Surprisingly, STIM1Δβ mice demonstrated a sexually dimorphic phenotype, where male STIM1Δβ mice were unaffected and female STIM1Δβ mice exhibited glucose intolerance, reduced insulin secretion, decreased β cell mass, increased α cell mass, and loss of key markers of β cell maturity and identity. RNA sequencing analysis showed downregulation of G protein-coupled estrogen receptor 1 (GPER1/GPR30), a non-canonical estrogen receptor, in HFD fed STIM1Δβ female islets. Consistent with this, STIM1 deficient β cells had blunted cAMP responses to stimulation with estradiol (E2) or a GPER1 agonist, whereas GPER1 knockdown or inhibition in β cell lines reduced the expression of markers of β cell maturity and function. Together, our results demonstrate a novel pathway in which STIM1-mediated SOCE orchestrates the maintenance of pancreatic β cell function and identity via estrogen signaling through GPER1 signaling.

## Results

### β cell-specific STIM1 deficiency does not alter systemic glucose tolerance at 8 weeks of age

To define the impact of STIM1 loss on *in vivo* pancreatic β cell function, a mouse model of β cell-specific STIM1 knockout (STIM1Δβ mice) was generated by crossing STIM1^fl/fl^ mice (Oh-Hora et al., 2008) with mice constitutively expressing *Ins1*-cre (Thorens et al., 2015). Appropriate genetic recombination was confirmed via PCR analysis (Figures S1A and S1B). Additionally, immunoblot performed in islets isolated from control and STIM1Δβ mice demonstrated efficient deletion of STIM1 protein (Figure 1A). Importantly, islets from STIM1Δβ mice did not show compensatory changes in mRNA expression of the two main P-type ATPases responsible for β cell ER Ca^2+^ uptake, *Serca2* and *Serca3*, or other SOCE molecular components, including *Stim2, Orai1*, or *Orai2* (Figure S1C).

**Figure 1.**
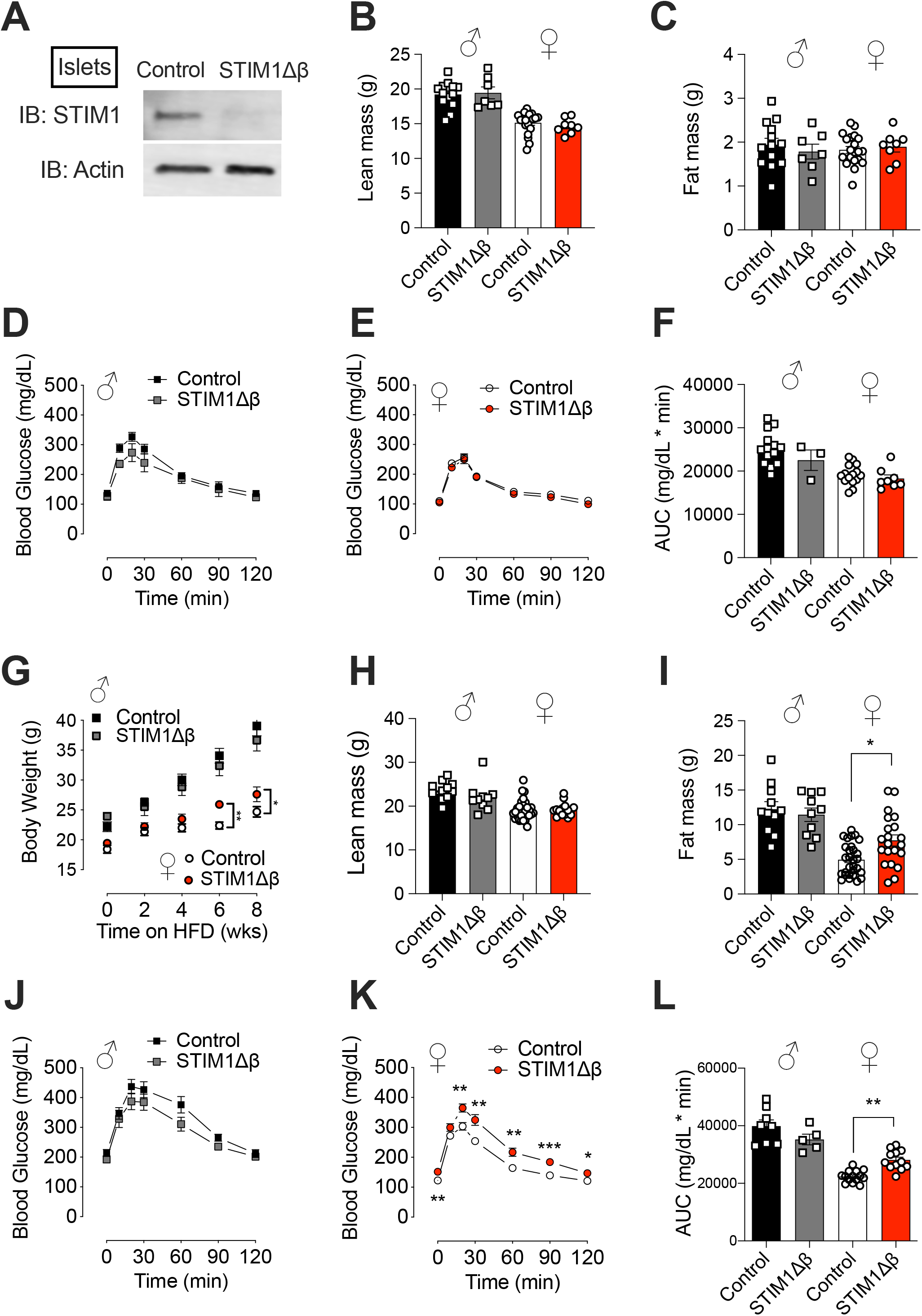
β cell-specific STIM1 deficiency causes glucose intolerance in female mice upon HFD feeding. (A-F) Control (STIM1fl/fl and unfloxed cre+) and STIM1Δβ mice were analyzed at 8 weeks of age. (A) Immunoblot analysis of islets isolated from control and STIM1Δβ mice using anti-STIM1 and anti-actin antibodies. (B and C) Lean (B) and fat (C) mass were measured using the EchoMRI 500 Body Composition Analyzer. (D-F) A GTT was performed (1.5 g/kg glucose dosed to lean mass) in male (D) and female (E) mice. (F) GTT results were analyzed using area under curve (AUC) analysis. (G-L) Control and STIM1Δβ mice were fed a HFD containing 60% of kcals from fat for 8 weeks, beginning at 8 weeks of age. (G) Changes in body weight in male and female mice were monitored over the HFD feeding period. (H and I) After HFD feeding for 8 weeks, lean (H) and fat (I) mass were measured using the EchoMRI 500 Body Composition Analyzer. (J-L) A GTT was performed (1.5 g/kg glucose dosed to lean mass) in male (J) and female (K) HFD-fed mice. (L) AUC analysis is shown graphically. Replicates are indicated by circles or squares; n ≥5 in each group. Results are displayed as mean ± SEM. Indicated differences are statistically significant: *p < 0.05; **p < 0.01; ***p < 0.001.

At 8 weeks of age, lean and fat mass were identical between control and STIM1Δβ mice in both male and female cohorts (Figures 1B and 1C). An intraperitoneal glucose tolerance test (GTT) showed no difference in systemic glucose homeostasis between control and STIM1Δβ mice in either males or females at this age (Figures 1D-1F).

### β cell-specific STIM1 deficiency increases weight gain and fat mass in HFD-fed female mice

Beginning at 8 weeks of age, male and female STIM1Δβ mice and littermate controls were fed HFD, in which 60% of calories were from fat. HFD led to weight gain in both genotypes and sexes. No differences in body weight were observed between male control and STIM1Δβ mice fed HFD for up to 8 weeks of diet treatment (Figure 1G). However, at 6 and 8 weeks of HFD feeding, female STIM1Δβ mice had increased body weight compared to female control mice (Figure 1G). After 8 weeks of HFD, there was no difference in lean mass between STIM1Δβ mice and their respective controls in either males or females (Figure 1H). However, female HFD-fed STIM1Δβ mice had higher fat mass compared to their HFD-fed female littermate controls (Figure 1I).

### β cell-specific STIM1 deficiency leads to glucose intolerance in female HFD-fed mice

After 8 weeks of HFD, control and STIM1Δβ mice of both sexes were subjected to a glucose tolerance test (GTT), in which the dose of glucose injected was based upon lean mass. Glucose tolerance was identical in male HFD-fed control and STIM1Δβ mice. In contrast, female HFD-fed STIM1Δβ mice had increased glucose excursions and worsened glucose tolerance when compared to control HFD-fed female mice (Figure 1J-1L). To determine whether β cell STIM1 ablation affected peripheral insulin sensitivity, an insulin tolerance test was performed in male and female control and STIM1Δβ mice after 10-weeks of HFD. No differences in insulin tolerance were detected between male HFD-fed control and STIM1Δβ mice or between female HFD-fed control and STIM1Δβ mice (Figure S2A-S2C).

Given the small but significant difference in body weight and composition between female HFD-fed control and STIM1Δβ mice, metabolic cage analysis was performed. No differences in food intake, respiratory exchange rate, or total activity counts were seen between these two groups of mice (Figures S2D-S2H). Together, these findings demonstrate a sexually dimorphic impact of β cell STIM1 deletion on systemic glucose metabolism in response to diet-induced obesity. The absence of insulin intolerance in HFD-fed female STIM1Δβ mice suggests that β cell dysfunction may underlie the phenotype.

### β cell-specific STIM1 deficiency decreases β cell function and mass in HFD-fed female mice

To assess β cell function, HFD-fed female control and STIM1Δβ mice were fasted for 6 hours, and serum insulin levels were measured 15 and 30 minutes after intraperitoneal injection of glucose (2 g/kg of lean mass). At 15 minutes after glucose injection, the increase in circulating insulin was significantly lower in female STIM1Δβ mice than in control mice (Figure 2A). In addition, fed insulin levels were significantly reduced in STIM1Δβ females (Figure 2B).

**Figure 2.**
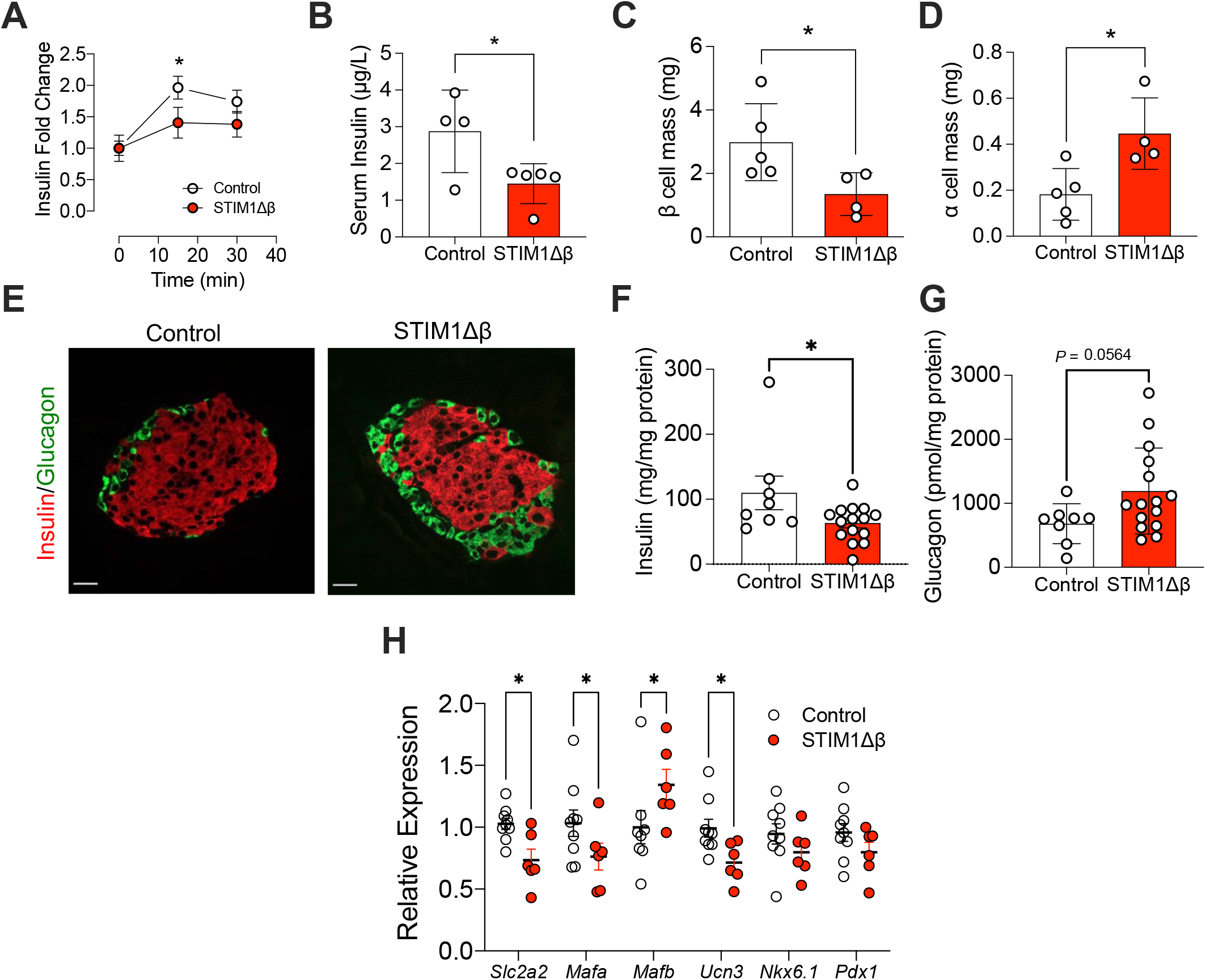
β cell-specific STIM1 deletion in female mice impairs insulin secretion due to a reduction in β cell mass and an increase in α cell mass. (A) After a 6 h fast, serum insulin levels were measured in control and STIM1Δβ female mice fed with HFD for 8 weeks at time 0 and 15 and 30 minutes after an intraperitoneal injection of glucose (2 g/kg glucose dosed to lean mass; n ≥ 5 per group). (B) Random fed serum insulin levels in HFD-fed control and STIM1Δβ female mice were determined by ELISA. (C and D) The total mass of insulin-positive β cells and glucagon-positive α cells within pancreata was quantified in paraffin tissue sections by immunostaining for insulin and glucagon in female mice fed with HFD for 8 weeks. (E) Representative images of islets in HFD-fed control and STIM1Δβ female mice; sections were stained with anti-insulin and anti-glucagon antibodies. (F and G) Islets from animals treated with HFD for 8 weeks were isolated and lysed, and intra-islet insulin (F) and glucagon (G) protein levels were measured by ELISA. (H) Islets isolated from HFD-fed control and STIM1Δβ female mice were analyzed by RT-qPCR for *slc2a2, mafa, mafb, unc3, nkx6*.*1*, and *pdx1*, normalized to *actb* levels. Replicates are indicated by circles; n ≥ 4 in each group. Indicated differences are statistically significant: *p < 0.05.

To test whether the decrease in circulating insulin concentration was related to changes in endocrine cell composition within the pancreas, β and α cell mass were quantitated in pancreatic sections from female control and STIM1Δβ mice following 8 weeks of HFD feeding. There was a significant reduction in β cell mass and a concomitant increase in α cell mass in female STIM1Δβ mice compared to controls (Figure 2C-2E). Consistently, analysis of pancreatic islets isolated from HFD-fed STIM1Δβ females revealed a significant reduction of insulin content and a trend towards an increase in glucagon levels (p=0.056, Figure 2F-2G). Given the observed changes in islet endocrine cell composition, we next measured the expression of a panel of genes associated with β cell function and maturity in islets isolated from female HFD-fed control and STIM1Δβ mice. This analysis revealed significantly reduced expression of markers of β cell identity (*Slc2a2, Mafa*, and *Ucn3* mRNA), whereas the expression of *Mafb*, a marker of α cell identity, was significantly increased. No significant differences were found in *Nkx6*.*1* or *Pdx1* levels (Figure 2H).

### STIM1-deficient β cells demonstrate loss of cellular identity

To determine which proteins are altered by STIM1 deletion in β cells, unbiased tandem mass tag (TMT) proteomics was performed in WT and STIM1KO INS-1 832/13 β cell lines (Kono et al., 2018). We identified a total of 6142 proteins in WT and STIM1KO INS-1 cells. Applying a cutoff of *P*<0.05 and a Benjamini-Hochberg post-test with a false discovery rate (FDR) <0.05, we identified 141 proteins that were differentially expressed between WT and STIM1KO cells. Furthermore, Metascape analysis with an enrichment value of at least 1.3 and p<0.05 showed that 40 pathways were differentially regulated between WT and STIM1KO cells (Zhou et al., 2019). Figure 3A shows the top 29 representative pathways, which includes terms such as “insulin signaling”, “glucagon signaling”, and “estrogen signaling”. The most significantly changed proteins in each of these pathways are shown as a heat map in Figure 3B. Notably, insulin-1 and insulin-2 expression were downregulated, while glucagon was upregulated in STIM1KO cells (Figure 3B). These results were confirmed by Western blotting, which showed reduced insulin and increased glucagon levels in STIM1KO compared to WT INS-1 cells (Figure 3C). For this analysis, α-TC cells were used as a positive control for the glucagon immunoblot.

**Figure 3.**
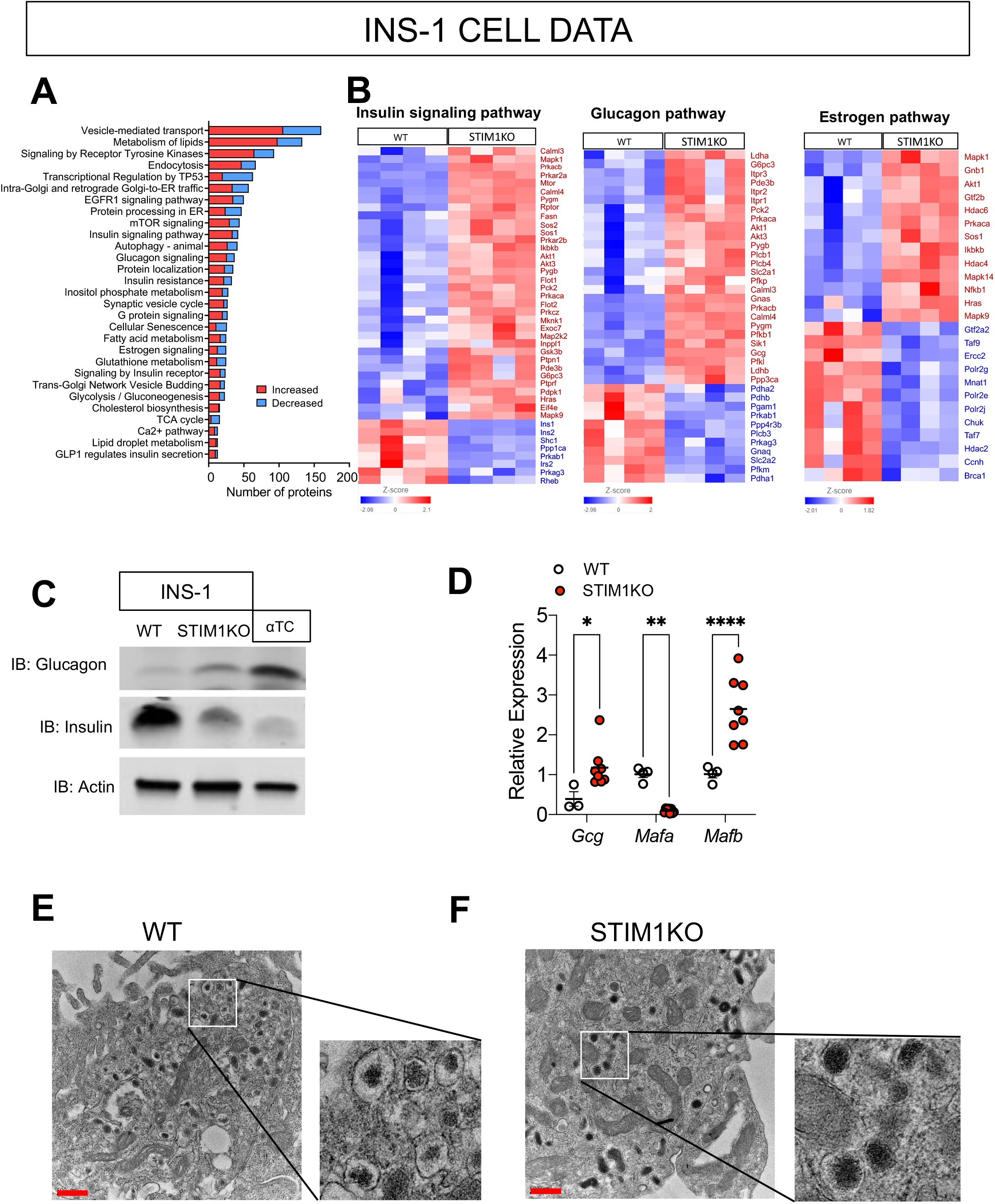
STIM1-deficient β cells demonstrate loss of maturity and identity. (A and B) WT and STIM1-deficient INS-1 832/13 cells (STIM1KO) were subjected to tandem mass tag MS/MS proteomics analysis. (A) Pathway analysis was performed using Metascape; 29 representative pathways/terms are illustrated. Red and blue represent the number of increased and decreased proteins in each pathway, respectively. (B) Abundance values of differentially expressed proteins in insulin signaling pathway, glucagon pathway, and estrogen pathway are represented in heatmaps. (C) Immunoblot analysis for glucagon, insulin, and actin in WT and STIM1KO INS-1 cells and α-TC cells. (D) RT-qPCR for *gcg, mafa*, and *mafb*, normalized to *actb* levels, in WT and STIM1KO INS-1 cells. (E and F) WT and STIM1KO cells were analyzed by electron microscopy. Representative images of insulin granule morphology from WT (E) and STIM1KO (F) INS-1 cells are shown (scale bar = 500 nm). Replicates are indicated by circles; n ≥ 4 in each group. Indicated differences are statistically significant; *p < 0.05, **p < 0.01, ****p < 0.0001 by one-way ANOVA (D).

Similar to islets isolated from STIM1Δβ mice, STIM1KO INS-1 cells demonstrated an mRNA signature of reduced β cell maturity and identity, i.e., upregulated expression of *gcg* and *mafb* mRNA and reduced expression of *mafa* mRNA (Figure 3D). Next, differences in the morphology of insulin granules between WT and STIM1KO INS-1 cells were evaluated by electron microscopy. WT INS-1 cells contained insulin granules with a typical morphology characterized by a dense homogeneous core and surrounding clear halo (Fava et al., 2012) (Figure 3E). In contrast, STIM1KO cells had densely filled granules without the surrounding halo, an appearance more typical of glucagon-containing granules (Brereton et al., 2014; Powers et al., 1990) (Figure 3F). In summary, unbiased and targeted analysis of STIM1KO INS-1 cell lines shows a phenotype similar to that observed in HFD-fed female STIM1Δβ mice—loss of β cell identity and function.

### RNA-seq analysis revealed alterations in estrogen-regulated pathways in islets from female STIM1Δβ mice

To obtain further mechanistic insight into the pathways leading to β cell dysfunction in female STIM1Δβ mice, islets were isolated from HFD-fed female control and STIM1Δβ mice and subjected to RNA sequencing analysis. Principal component analysis showed a clear separation between control and STIM1Δβ mice (Figure 4A). We identified 836 differentially expressed genes based on a threshold fold-change ≥1.5 and FDR <0.05. Of these, 308 genes were downregulated, and 528 genes were upregulated (Figure S3A). The expression pattern of the top 50 differentially expressed genes (25 upregulated and 25 downregulated) is shown in Figure 4B. Pathway analysis of the RNA sequencing data identified 55 pathways that were significantly modulated (*P* < 0.05), and 30 representative pathways are listed in Figure 4C. Several pathways relevant to β cell function and the pathogenesis of T2D were identified, including “mitochondrial dysfunction”, “cholesterol biosynthesis”, “endocytosis”, and “phagocytosis” (Figure 4C). Notably, estrogen receptor signaling was also identified as a significantly modulated pathway (Figure 4C). This finding was interesting, as we observed a sexually dimorphic metabolic phenotype in the STIM1Δβ mice. Therefore, we utilized IPA to search for potential upstream regulators of differentially expressed genes, focusing on estrogen and estrogen-related molecules and their corresponding downstream targets (Figure S3B). Twenty representative gene ontology terms obtained by the functional enrichment analysis of the identified estrogen-regulated genes are shown in Figure S3C: they include “glucagon signaling”, “mitochondrial biogenesis”, “G-protein coupled receptor signaling”, and “glucose metabolism”. We reasoned that this novel link between STIM1 and estrogen receptor signaling could contribute to the sexually dimorphic phenotype observed in our STIM1Δβ mice.

**Figure 4.**
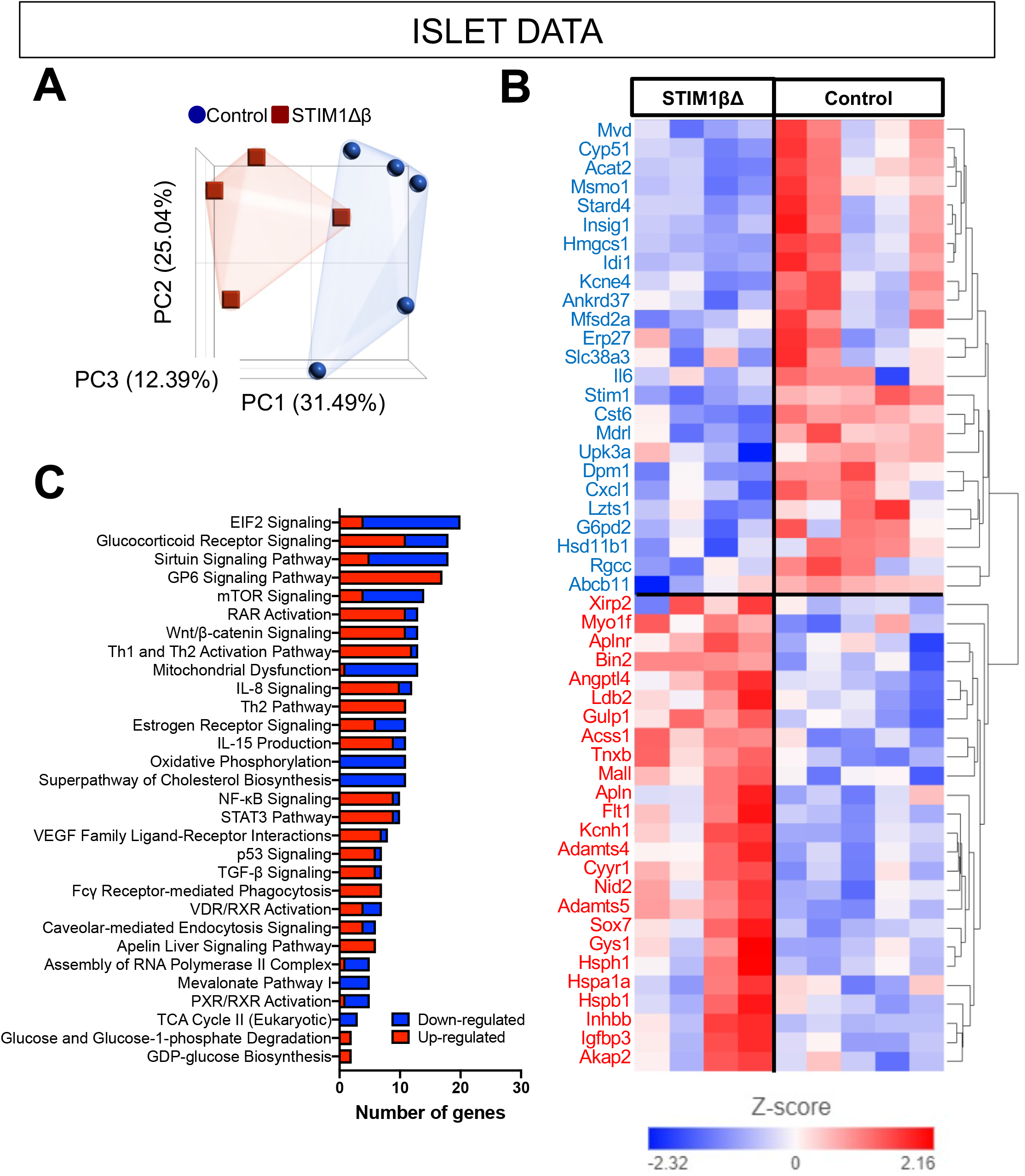
RNA-seq analysis revealed differential expression of genes related to pancreatic β cell dedifferentiation in islets from female STIM1Δβ mice. (A-C) Pancreatic islets isolated from HFD-fed control and STIM1Δβ female mice were subjected to RNA-sequencing analysis using 100 ng of total RNA per sample. (A) Principal component analysis revealed clear separation between control and STIM1Δβ islets. Log cpm values were used to generate the PCA plot. (B) Top 50 (25 upregulated and 25 downregulated) differentially expressed genes are shown in a heat map. Genes in blue indicate downregulated genes in STIM1Δβ islets and genes in red indicate upregulated genes in STIM1Δβ islets. (C) Functional enrichment analysis was performed using IPA; 20 representative pathways are illustrated. Red and blue indicate the number of up and downregulated genes in each pathway, respectively.

### STIM1 deletion or inhibition of SOCE downregulates GPER1 expression and signaling in WT mouse islets and INS-1 cells

Because RNA sequencing showed differential regulation of estrogen-related pathways, we next analyzed the data to determine the expression pattern of the three main estrogen receptors (ER) (Mauvais-Jarvis et al., 2017). The expression of ERα and β was unaffected (Figure S3D), while GPER1, a G protein-coupled receptor that is an important regulator of 17-β estradiol (E2) signaling, was downregulated in STIM1Δβ islets (Figure 5A). This reduction in GPER1 was confirmed at the mRNA and protein level by RT-qPCR and immunoblot analysis of islets from a separate cohort of female control and STIM1Δβ mice (Figures 5B-5D). To further clarify the relationship between the regulation of ER Ca^2+^ levels and GPER1 expression, islets from WT C57Bl/6J female mice were treated with SOCE inhibitors (ML-9, 2-APB, or AncoA4) for 24 hours. All three inhibitors reduced the expression of GPER1 mRNA (Figure 5E), indicating that SOCE is required for the maintenance of GPER1 expression in female islets.

**Figure 5.**
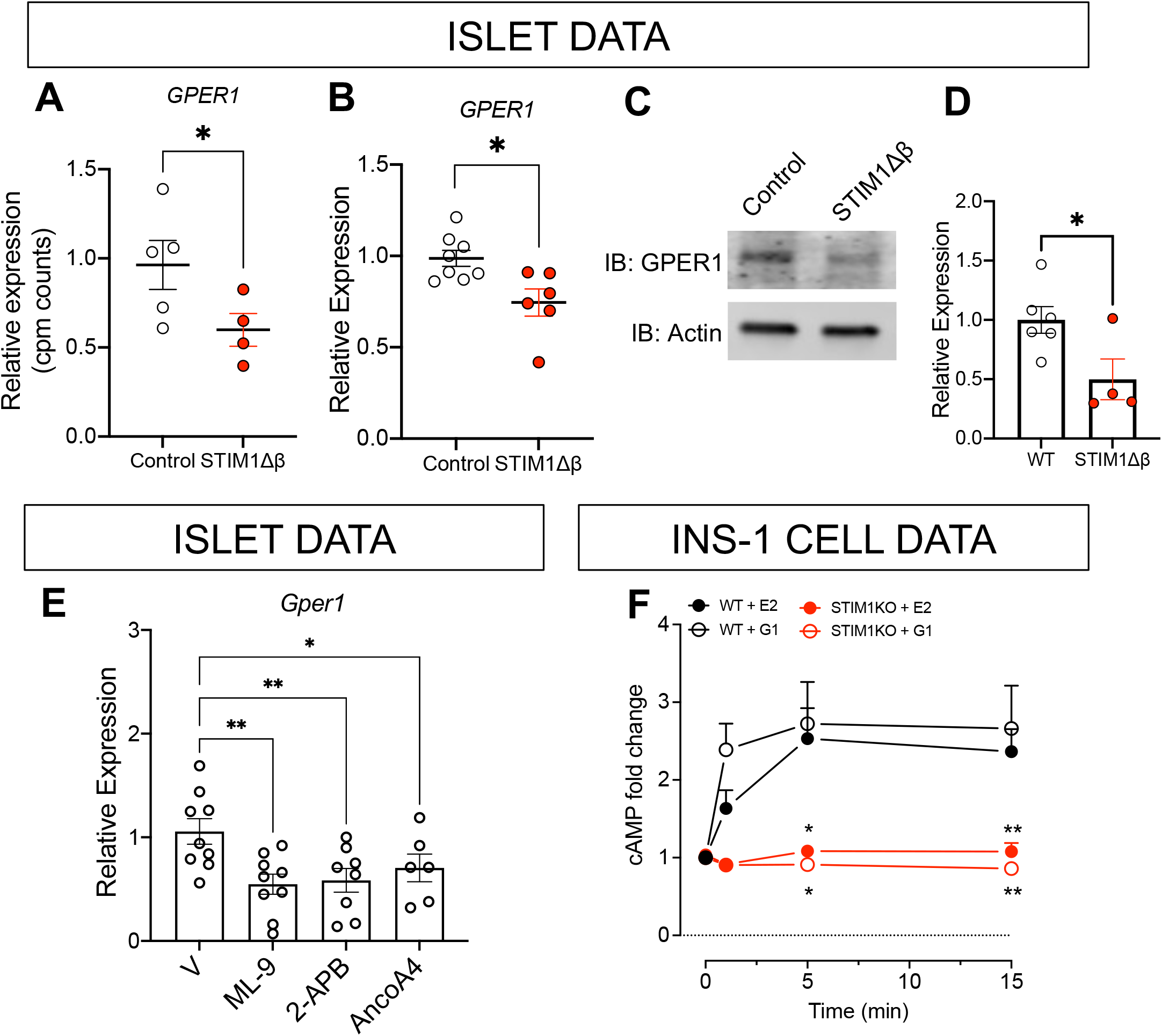
Inhibition of SOCE decreases expression of GPER1 in pancreatic islets and INS-1 cells. (A) Cpm counts of *gper1* obtained from RNA-seq analysis were compared between HFD-fed control and STIMΔβ female mouse. (B) Expression levels of *gper1* between female control and STIM1Δβ islets were compared by RT-qPCR normalized to *actb*. (C and D) Immunoblot of GPER1 and actin from islets of HFD-fed control and STIM1Δβ female mice (C) and quantitation of GPER1 after normalization to actin (D). (E) Islets isolated from 8-week-old C57Bl/6J female mice were treated with ML9, 2APB, or AncoA4, and *gper1* expression level was determined by RT-qPCR normalized to *actb*. (F) Intracellular cAMP mobilization was measured using an HTRF assay. WT or STIM1KO INS-1 cells were treated with E2 (10 nM) or G1 (10 nM) for 1, 5, and 15 min. The data were normalized to the baseline of the respective genotype (WT or STIM1KO). Replicates are indicated by circles; n≥4 in each group. Indicated differences are statistically significant; *p < 0.05, **p < 0.01.

To directly test the relationship between E2 signaling and STIM1, WT and STIM1KO cells were treated with 10 nM E2 or G-1, an agonist of GPER1 (Sharma et al., 2020). Treatment with E2 or G-1 for 15 minutes resulted in an approximately 2.5-fold increase in intracellular cAMP concentration in WT INS-1 cells, indicating effective activation of this PKA-associated GPCR (Figure 5F). However, the level of cAMP in STIM1KO cells remained essentially unchanged during E2 or G-1 treatment (Figure 5F), confirming a defect in estrogen signaling in the absence of STIM1.

### GPER1 downregulation or inhibition in INS1 cells decreases expression of β cell identity genes and increases expression of glucagon

Finally, to test whether GPER1 is required for the maintenance of β cell identity, INS-1 832-13 cells were transfected with siRNA targeted to *Gper1*. Efficient knockdown of *Gper1* was confirmed by RT-qPCR (Figure 6A). Markers of β cell identity, including *Slc2a2, Mafb*, and *Ucn3*, were significantly reduced in the *Gper1* knockdown cells. Furthermore, knockdown of *Gper1* increased *Gcg* gene expression in this β cell line (Figure 6A). Treatment of INS-1 cells with E2 or with the GPER1 antagonist G-15 confirmed the changes in gene expression observed via siRNA knockdown; there was a significant decrease in the β cell identity genes *Slc2a2* and *Ucn3* and a significant increase in *Gcg* expression (Figures 6B-6F).

**Figure 6.**
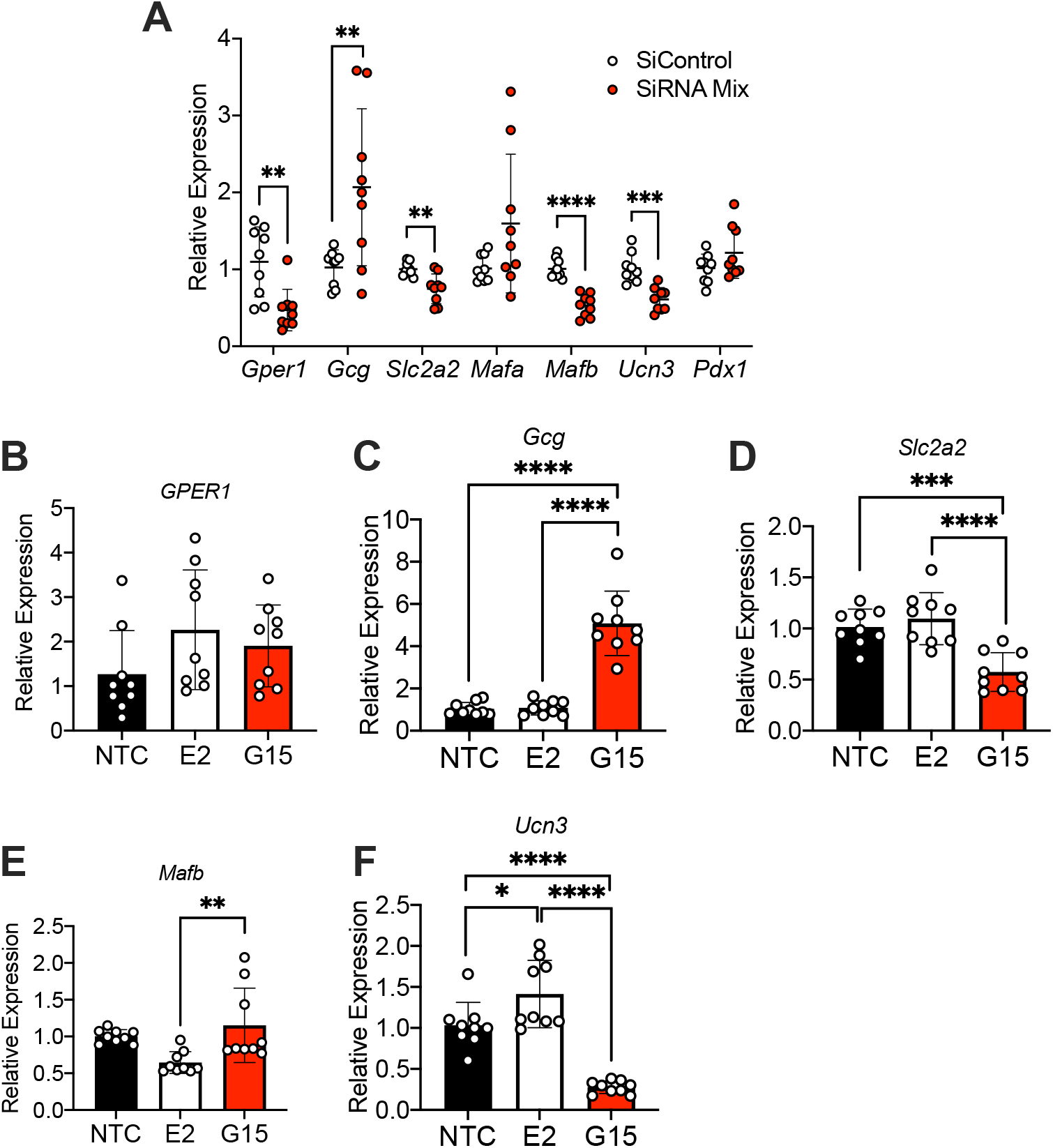
GPER1 knockdown and inhibition in INS-1 cells leads to a loss of β cell identity and increased glucagon expression. (A) INS-1 832-13 cells were transfected with siRNA targeted to *gper1*. Expression of *gper1, gcg, glut2, mafa, mafb, unc3*, and *pdx1* was measured by RT-qPCR and normalized to *actb*. (B-F) INS-1 832-13 cells were treated with the GPER1 agonist E2 or the GPER1 antagonist G-15. Expression of *gper1, gcg, glut2, mafb, and unc3* was measured by RT-qPCR and normalized to *actb*. Replicates are indicated by circles; n≥9 in each group. Indicated differences are statistically significant; * p < 0.05, ** p < 0.01, *** p < 0.001, **** p < 0.0001.

## Discussion

Ca^2+^ plays a vital role in normal β cell function by regulating critical steps in insulin production, maturation, and secretion. The fidelity of many of these processes depends on the maintenance of distinct Ca^2+^ stores, organized at the cellular and organelle level. The ER serves as the dominant intracellular Ca^2+^ store, and ER Ca^2+^ depletion represents a common pathway in response to multiple metabolic stressors that contribute to diabetes pathophysiology, including pro-inflammatory cytokines, high glucose, and lipotoxicity (Kono et al., 2012; Lytrivi et al., 2020; Shyr et al., 2019; Yamamoto et al., 2019). Under physiological conditions, reductions in ER Ca^2+^ trigger a tightly regulated rescue mechanism known as SOCE or Ca^2+^ release-activated Ca^2+^ current. Genetic deletion or pharmacological inhibition of SOCE or dominant-negative mutants of Orai1 or TRPC1 reduce glucose- and GPR40-mediated insulin secretion in rat islets and clonal β cell lines (Kono et al., 2018; Sabourin et al., 2015; Usui et al., 2019). Interestingly, STIM1, a key component of SOCE, also interacts with and regulates the activity of the sulfonylurea receptor 1 (SUR1) subunit of the KATP channel in β cells (Leech et al., 2017), suggesting a role in insulin secretion.

We have shown that levels of STIM1 are decreased in islets from donors with T2D, and that STIM1 loss is linked with β cell ER stress (Kono et al., 2018). However, the *in vivo* role of STIM1 in the pathophysiology of T2D is not well established. Therefore, in this study we generated mice with a β cell-specific STIM1 deletion (STIM1Δβ mice) using cre-loxP recombination. We demonstrated that STIM1 deficiency in β cells causes glucose intolerance and β cell dysfunction in female, but not male, mice upon HFD feeding. Further characterization of female STIM1Δβ mice revealed that deletion of STIM1 impaired insulin secretion as a result of reduced β cell mass and increased α cell mass. RT-qPCR analyses of islets from HFD-fed control and STIM1Δβ female mice demonstrated that STIM1 deficiency resulted in the downregulation of key markers of β cell maturity, such as *Slc2a2* (GLUT2), *Mafa*, and *Ucn3*, and an increase in *Mafb*, an α Cell marker, suggesting that STIM1 is critical for the maintenance of β cell identity, maturity, and function in female mice.

While the sexually dimorphic pattern of glucose intolerance and β cell dysfunction in our STIM1Δβ mice was unexpected, the preferential loss of β cell identity in female mice has been seen in other studies. In one of the first reports to describe β cell dedifferentiation as a component of diabetes pathophysiology, multiparous FoxO1 knockout female mice, but not male mice, had a reduction of β cell mass and an increase in α cell mass, with reduced expression of *Pdx1, Nkx6*.*1, MafA, Pcsk2, Gck*, and *Glut2* (Talchai et al., 2012). In addition, Zhang and collaborators demonstrated that β cells from female Sprague Dawley rats undergo dedifferentiation when challenged with long-term fluctuating glycemia and these changes were associated with oxidative stress (Zhang et al., 2019).

Analysis of our RNA-sequencing dataset from islets of female mice showed alterations in estrogen-related signaling pathways, raising the possibility of an underlying mechanistic connection between STIM1, β cell differentiation/function, and sex steroids. Indeed, studies in humans and rodents have shown that an abrupt disruption of the ovarian hormonal axis via oophorectomy leads to glucose intolerance and diabetes (Appiah et al., 2014; Santos et al., 2016). Some, but not all, studies have suggested a similar response during natural menopause, but this remains somewhat controversial (Brand et al., 2015; Kim et al., 2011; Ren et al., 2019). However, 17β-estradiol (E2) signaling in rodent and human β cells is known to play a protective role against diabetic stressors, thus supporting islet survival, biosynthesis, lipid homeostasis, and proliferation (Alonso-Magdalena et al., 2008; Le May et al., 2006; Mauvais-Jarvis et al., 2017; Yamabe et al., 2010). Three estrogen receptors have been identified in the β cell: estrogen receptor α, estrogen receptor β, and GPER1 (Mauvais-Jarvis et al., 2017). Interestingly, we found selective downregulation of GPER1 expression in STIM1Δβ female mouse islets, with preserved expression of ERα and β.

GPER1, also known as GPR30, belongs to a class of G protein-coupled receptors and functions as an estrogen receptor. Several studies link GPER1 signaling to β cell function. Islets isolated from GPER1-deficient mice demonstrate reduced GSIS (Martensson et al., 2009), and GPER1-deficient female mice are more susceptible to STZ-induced diabetes (Liu et al., 2009). Further, GPER1 impacts β cell proliferation via the activity of miR-338-3p (Jacovetti et al., 2012). Lombardi and coworkers also established the protective effect of E2 signaling in dedifferentiation of β cells by showing that the pre-treatment of islets with E2 rescued GlcN-induced β cell dedifferentiation, and this effect was reversed by the GPER1 antagonist G-15 (Lombardi et al., 2012).

Our studies now show that GPER1 requires STIM1 to protect β cells from diabetic stressors. First, we measured cAMP mobilization, a readout for GPER1 activity (Filardo, 2002; Filardo et al., 2002), in WT and STIM1 KO INS-1 cells. As expected, cAMP levels in WT cells were significantly increased within 5 minutes of E2 and G-1 (GPER1 agonist) treatment. However, this cAMP response was ablated in STIM1KO cells, suggesting that STIM1 is required for E2 signaling through GPER1 in β cells. Second, we showed that siRNA-mediated knockdown of GPER1 or treatment with a GPER1 antagonist (G-15) decreased the expression of markers of β cell identity in INS-1 cells. Lastly, we showed that inhibition of SOCE decreased islet expression of GPER1, suggesting that STIM1 also regulates estrogen signaling. Therefore, our experiments established that STIM1 is required for GPER1-mediated protection of β cells and maintenance of β cell identify.

There are some limitations of our study that should be acknowledged. First, the INS-1 832/13 cell line (RRID: CVCL_7226) was originally isolated from a male rat (Hohmeier et al., 2000), and we observed a female-specific phenotype in our mouse model. However, we linked this sexual dimorphism to GPER1 signaling, and we have shown that GPER1 is expressed in INS-1 cells. Therefore, this cell line was an appropriate model for our *in vitro* studies. Further studies will be needed to determine the applicability of these findings to human models: GPER1 is expressed in human pancreatic islets; however, the relationship between STIM1, GPER1 expression, and estrogen signaling in humans remains to be demonstrated.

Notwithstanding these limitations, our findings show the importance of STIM1 in the maintenance of pancreatic β cell identity in a rodent model of diet-induced obesity. STIM1 deficiency in β cells showed an unexpected sexually dimorphic presentation of metabolic dysfunction and β cell dedifferentiation. This female-specific phenotype was due, in part, to downregulation of E2 signaling through GPER1 in the absence of STIM1. E2 contributes to the maintenance of β cell mass and identity by providing protection against stressors that cause β cell dedifferentiation. Therefore, the data obtained in this study reveal novel insights into sex differences in the development of T2D, and demonstrate the importance of STIM1 and Ca^2+^ signaling in the hormonal regulation of T2D etiology. Additionally, Orai inhibitors are currently being tested for efficacy in disease states including pancreatitis (Bruen et al., 2021). Our results suggest that efforts to restore STIM1 expression or increase activation of SOCE may be considered as potential therapeutic strategies for T2D in females.

## Supporting information

ARRIVE Checklist

Key Resources Table

## Acknowledgments

This work was supported by National Institute of Diabetes and Digestive and Kidney Diseases grants R01DK093954, R01DK127236, U01DK127786, R01DK127308, UC4DK104166 (to C.E.-M.), and 5F30DK123996-03 (to P.S.), U.S. Department of Veterans Affairs Merit Award I01BX001733 (to C.E.-M.), and gifts from the Sigma Beta Sorority, the Ball Brothers Foundation, the George and Frances Ball Foundation (to C.E.-M.). The funders had no role in study design, data collection and analysis, decision to publish, or preparation of the manuscript. The authors acknowledge the support of the Islet and Physiology and Translation Cores of the Indiana Diabetes Research Center (P30-DK-097512). The mass spectrometry experiments in this manuscript were performed in the Indiana University School of Medicine Center for Proteome Analysis. The authors would like to thank Drs. Emma H. Doud, Amber L. Mosley, Guihong Qi, and Jaison Arivalagan for their professional support on the mass spectrometry analysis. The authors would also like to thank Dr. Emily Anderson-Baucum for her helpful advice and edits, and Kara Orr, Lata Udari, and Jacqueline Aquino (Indiana University) for their technical assistance.

## Author Contributions

P.S. designed and conducted the experiments, performed data analysis, and wrote the manuscript; M.R.M conducted experiments, performed data analysis, and assisted with writing of the manuscript; P.K. performed bioinformatics analysis on RNA-sequencing data, interpreted data, and edited the manuscript; C.L. contributed to the design and execution of experiments, analyzed data, and assisted with the writing of the manuscript; T.K. contributed to funding acquisition, conception and design of the study, data analysis, interpretation, and manuscript writing/editing; C.E-M. directed the funding acquisition, designed the experiments, analyzed data, and wrote the manuscript. C.E-M. is the guarantor of this work, had full access to all of the study data, and takes responsibility for the integrity and accuracy of the data.

## Supplemental Figure Legends

**Figure S1.**
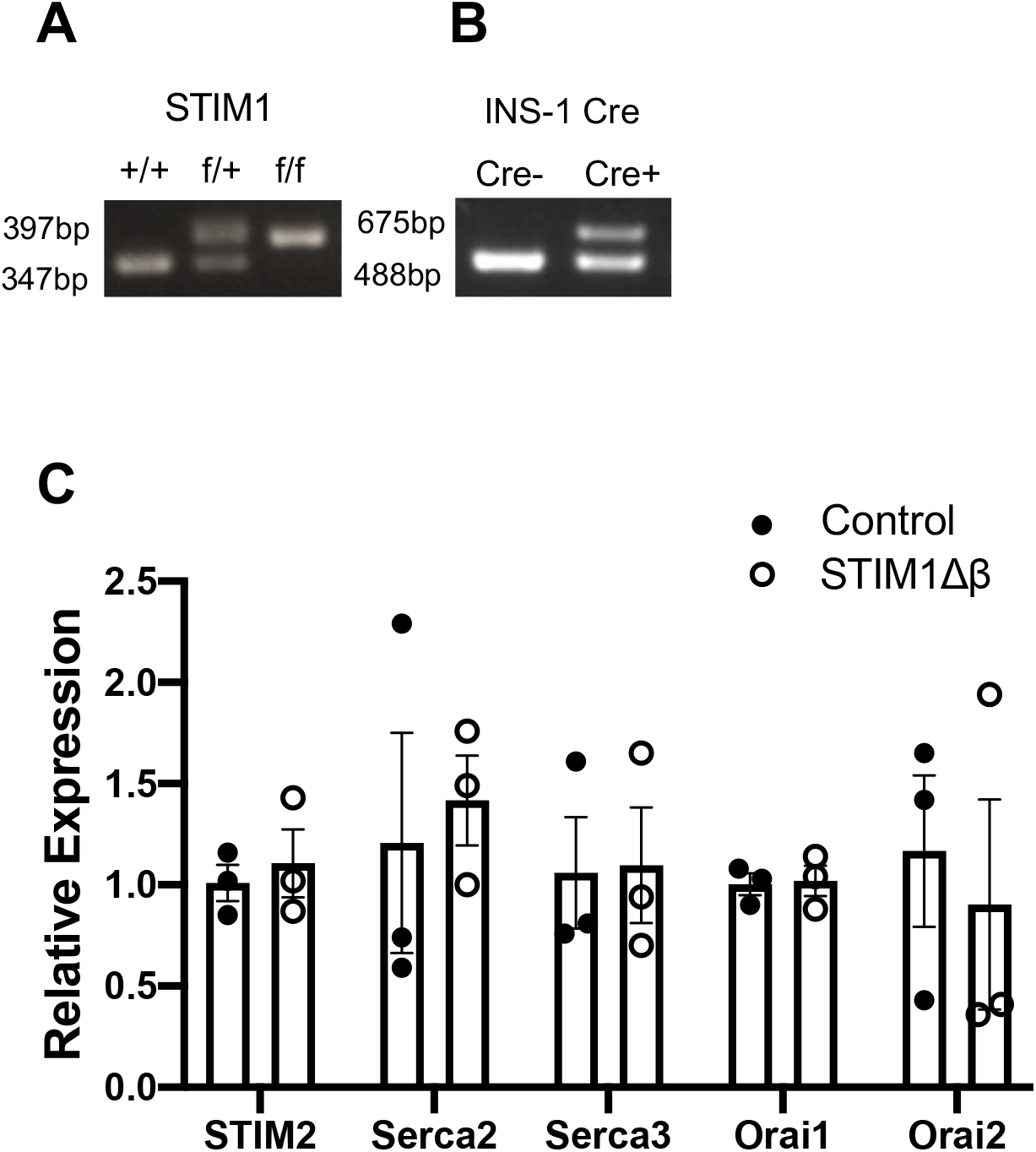
β cell-specific recombination of STIM1 does not alter expression of other components of SOCE. (A-B) Genotyping was performed according to protocols from Jackson Laboratories. Gel images of bands of PCR products from DNA from (A) STIM1fl/fl mice and (B) INS-1 cre mice. (C) RT-qPCR was performed using islets isolated from 8-week-old control and STIM1Δβ female mice (n=3 per group).

**Figure S2.**
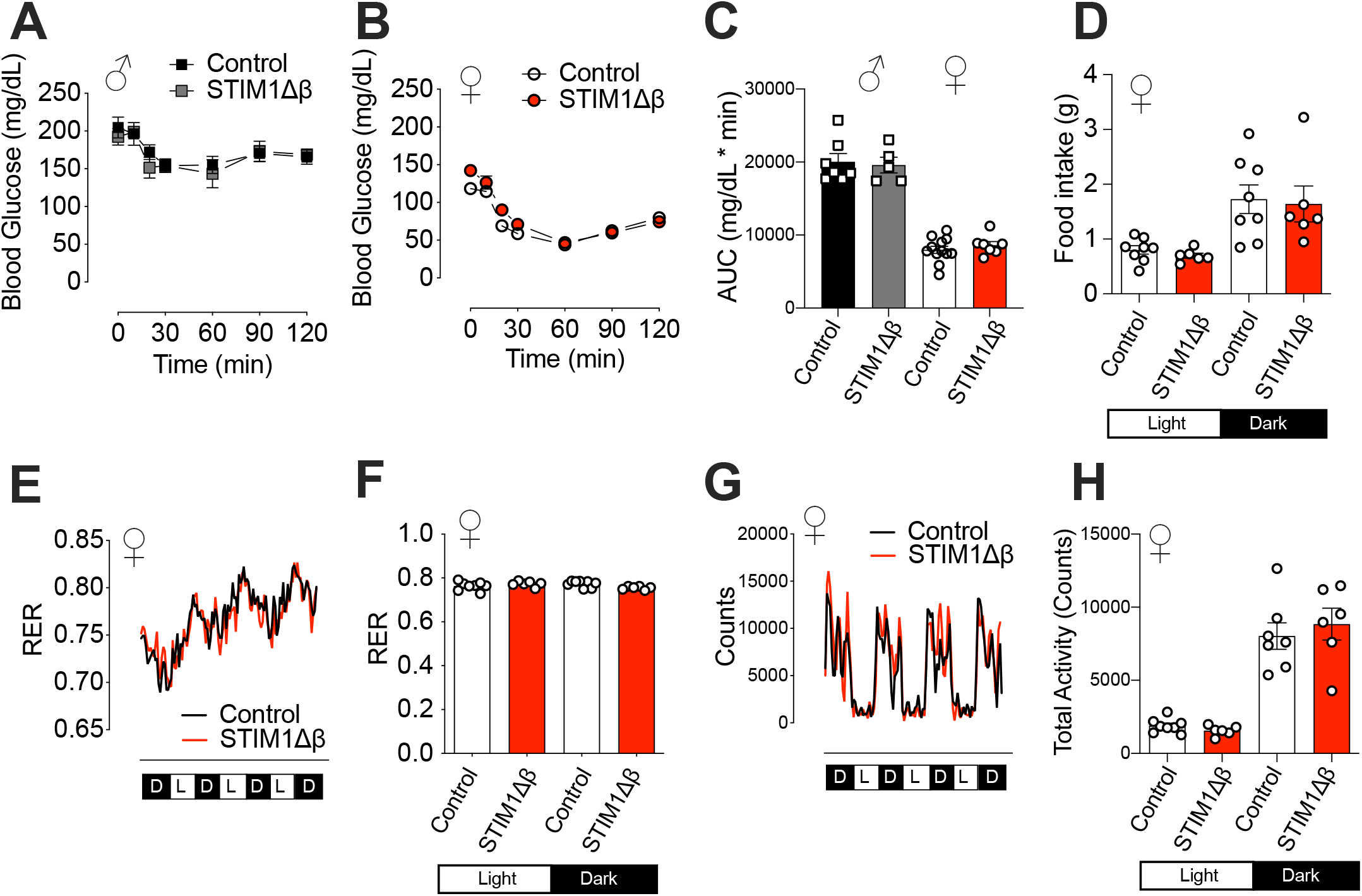
HFD-fed STIM1Δβ mice exhibit comparable insulin sensitivity, food intake, energy expenditure, and activity levels. (A-C) Insulin tolerance test (ITT; 0.75 mU/kg body weight) was performed in male (A) and female (B) mice after 10 weeks of HFD feeding following a 4 h fast. (C) Area under curve (AUC) quantitation. (D) After 10 weeks of HFD feeding, food intake by control and STIM1Δβ female mice was measured through the TSE LabMaster Metabolism Research Platform feeding monitor. (E-H) Energy expenditure was determined by metabolic cage testing, and was analyzed using the respiratory exchange ratio (E and F) and total activity (G and H). Results are displayed as mean ± SEM (n ≥ 6 per group).

**Figure S3.**
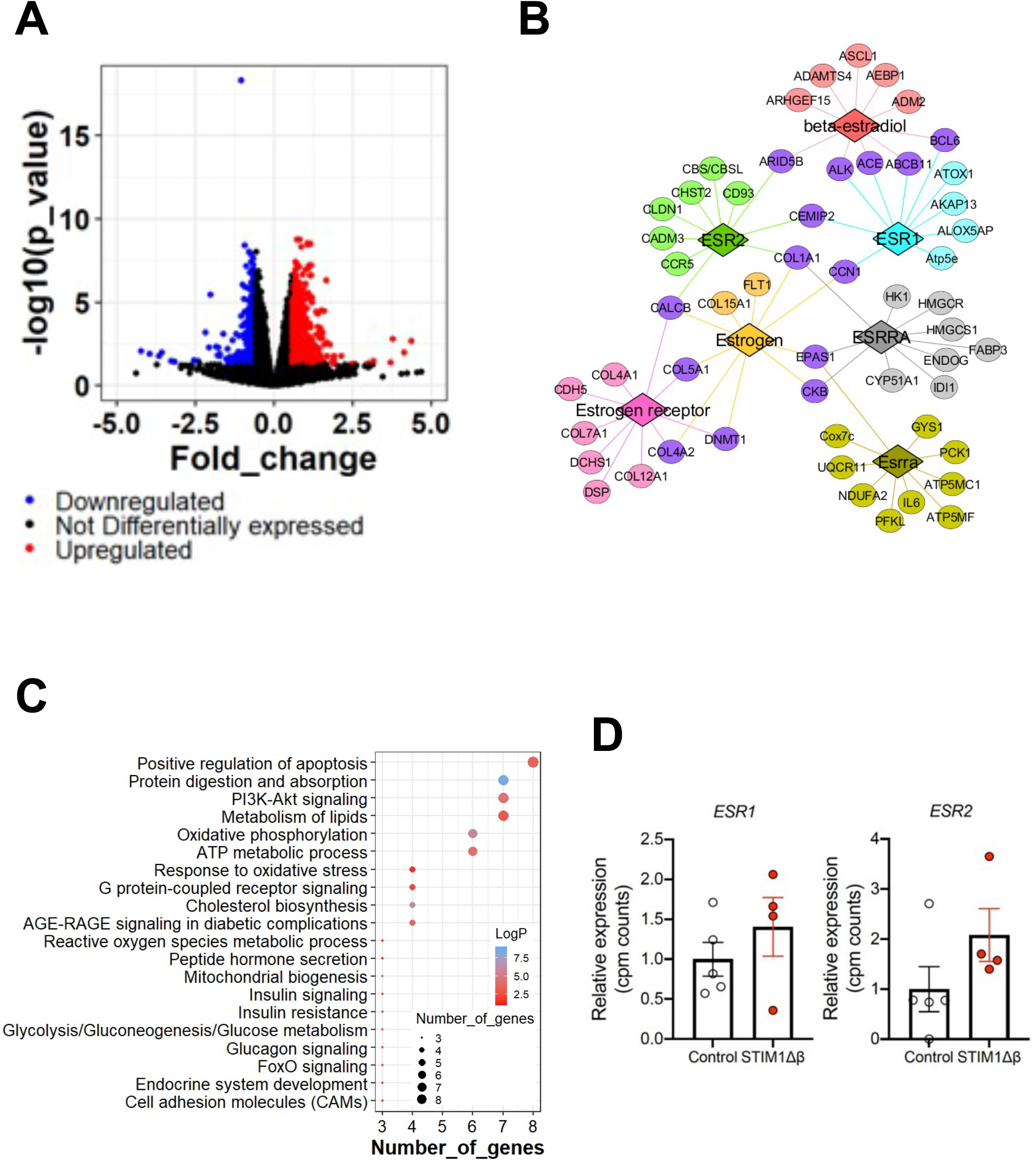
RNA sequencing data from islets. Data generated from RNA-seq analysis are represented in various plots. (A) Volcano plot showing differentially regulated genes with downregulated in blue and upregulated in red. (B) Upstream regulators were identified from IPA with a focus on estrogen and estrogen receptors. Each of the estrogen-associated upstream regulators (diamond shaped) are indicated with a unique color and the downstream regulated genes (circles) are indicated in the same color. Downstream genes common between any two or more upstream regulators are indicated in purple. (C) Dot plot depicts the functional enrichment analysis for estrogen-regulated downstream genes. (D) Relative expression of *Esr1* and *Esr2* genes measured as cpm counts: comparison between control and STIM1Δβ mice.

## STAR Methods

### RESOURCE AVAILABILITY

#### Lead Contact

Further information and requests for resources and reagents should be directed to and will be fulfilled by the lead contact, Carmella Evans-Molina.

#### Materials Availability

This study did not generate new unique reagents.

#### Data and Code Availability

The RNA sequencing data generated during this study are available at GEO repository, accession #GSE208135.

Raw and searched mass spectrometry data has been uploaded to MassIVE with a repository ID of MSV000089927.

### EXPERIMENTAL MODEL AND SUBJECT DETAILS

#### Animals

Male mice, strains B6(Cg)-Ins1^tm1.1(cre)Thor^/J (Ins1-Cre) and B6(Cg)-Stim1^tm1Rao^/J (STIM1^fl/fl^), were purchased from Jackson Laboratory, and colonies were established in our facility. Pancreatic β cell-specific STIM1 knockout (STIM1Δβ) mice were generated by crossing Ins1-Cre mice and STIM1^flox/flox^ mice. The STIM1^fl/fl^ and INS1-Cre strains were genotyped using Jackson Laboratory PCR protocols 25780 and 18406, respectively.

Mice were kept in sterilized Max 75 plastic cages (Alternative Design) under standard environmental temperature conditions (22 ± 2°C) and relative humidity (30-70%), and a 12:12 h light:dark cycle. All animal experiments were approved by the Indiana University Animal Care and Use Committees, were conducted in accordance with the NIH Guide for the Care and Use of Laboratory Animals, and were preformed according to ARRIVE (Animal Research: Reporting of In Vivo Experiments) guidelines (Percie du Sert et al., 2020). For metabolic phenotyping studies, male and female age-matched littermate controls were housed together in a sex-stratified manner with ≤5 mice per cage. Reported control values consist of results obtained from both INS1-Cre-negative STIM1^flox/flox^ (Cre-controls) and Cre positive STIM1^+/+^ (Cre+ controls lacking LoxP).

#### Cell lines

The STIM1 KO cell line (STIM1KO) was used as previously reported (Kono et al., 2018). STIM1KO and WT INS-1 832/13 cells (RRID: CVCL_7226) were maintained at 37°C in an atmosphere supplemented with 5% CO_2_ and grown in phenol red-free RPMI1640 medium (Gibco) containing 10% charcoal-stripped fetal bovine serum (Millipore Sigma), 100 U/mL penicillin, 100 μg/mL streptomycin (Invitrogen), 10 mM HEPES (Gibco), 2 mM L-glutamine (Sigma), 1 mM sodium pyruvate, and 50 μM β-mercaptoethanol (Sigma). The α-TC cell line was obtained from ATCC (alphaTC1 Clone 9, CRL-2350; RRID: CVCL_0150), and was cultured in low-glucose DMEM (Gibco) containing 10% charcoal-stripped fetal bovine serum, 100 U/mL penicillin, 100µg/mL streptomycin, 15 mM HEPES, and 0.1 mM L-glutamine.

### METHOD DETAILS

#### Metabolic studies

Mouse body weight was measured at indicated time points, and body composition (lean and fat mass) was determined using the EchoMRI-500 Body Composition Analyzer (EchoMRI; RRID:SCR_017104), as previously described (Sims et al., 2013). Mice were fed *ad libitum* with a high-fat diet (HFD, 60% kCal fat, Research Diets, catalog #D12492) starting at 8 weeks of age. After 8 weeks of HFD feeding, glucose tolerance tests (GTT) and insulin tolerance tests (ITT) were performed. GTT was performed after 6 hours of fasting followed by intraperitoneal (IP) administration of 1.5 g d-glucose/kg lean mass (Ayala et al., 2010). Insulin tolerance tests (ITT) were performed after 4 h of fasting followed by IP administration of 0.75 U insulin (Novo Nordisk) per kg body weight. Blood glucose levels were measured using a Contour glucometer (Bayer).

Glucose-stimulated insulin secretion (GSIS) was measured *in vivo* following 6 h of fasting. Serum was collected via the tail vein at 0, 15, and 30 min after an IP injection of glucose (2 g/kg lean body mass). Insulin levels in serum and tissues were measured using a mouse insulin ELISA kit (Mercodia).

#### Isolation of mouse islets

Mouse pancreatic islets were isolated by collagenase digestion as previously described (Stull et al., 2012). Male and female mouse islets were isolated at 8 weeks of age or after 8 weeks of HFD feeding. Islets were cultured in phenol red-free, low-glucose DMEM supplemented with 10% charcoal-stripped fetal bovine serum, 100 U/mL penicillin, 100 µg/mL streptomycin, and 2 mM L-glutamine.

#### Metabolic caging analysis

Metabolic caging analysis was performed as previously described (Sims et al., 2013). Briefly, indirect calorimetry measurements were performed after 8 weeks of HFD using the LabMaster Metabolism Research Platform (TSE Systems) equipped with a calorimeter, feeding monitor, and activity monitoring system. Metabolic parameters were measured after a 24 h acclimation period; data were collected every 10 min for 24 h. Respiratory quotient (RQ) and energy expenditure (EE) were calculated as previously described (Sims et al., 2013), and results are presented as mean measurements over a 24 h period as well as during light (0700–1900) and dark (1900–0700) cycles. Percent relative cumulative frequency (PRCF) curves were generated from RQ values as previously described (Sims et al., 2013).

#### Immunoblotting

Approximately 1.5 × 10^5^ INS-1 cells or 100–125 mouse islets were washed with PBS and lysed with buffer containing 50 mM Tris (pH 8.0), 150 mM NaCl, 0.05% deoxycholate, 0.1% IGEPAL CA-630, 0.1% SDS, 0.2% sarcosyl, 10% glycerol, 1 mM DTT, 1 mM EDTA, 10 mM NaF, EDTA-free protease inhibitors, phosphatase inhibitors, 2 mM MgCl_2_, and 0.05% v/v Benzonase Nuclease. Protein concentration was measured using the Bio-Rad DC protein assay (Bio-Rad) and a SpectraMax iD5 multi-mode microplate reader (Molecular Devices). Equal concentrations of proteins were loaded onto a 4-20% Mini-Protean TGX gel in a Mini-Protean Tetra apparatus (Bio-Rad). Proteins were transferred to a PVDF membrane and blocked with Intercept Blocking Buffer (LI-COR) prior to incubation with primary antibodies. Bound primary antibodies were detected by antibodies listed in the Key Resources table using indicated dilutions. Immunoblots were scanned with an LI-COR Odyssey 1828 scanner (LI-COR, RRID:SCR_014579) and analyzed densitometrically using Image Studio software (LI-COR, RRID:SCR_015795).

For immunoblotting of insulin and glucagon, a modified protocol of Okita and colleagues was used (Okita et al., 2017). Cells were lysed in an SDS sample buffer (75 mM Tris-HCl, pH 6.8, 1% SDS, 10% glycerol, 2.5% sucrose). After determining protein concentration using a BCA protein quantification assay (Bio-Rad) according to the manufacturer’s protocol, 2-mercaptoethanol (final concentration, 5%, Fisher Scientific) and bromophenol blue (final concentration, 0.00125%) were added, and the mixture was heated at 95°C for 5 min. An aliquot corresponding to 2 micrograms of protein was subjected to Western blotting. Mini-PROTEAN Tris/Tricine Precast Gels, 16.5% (Bio-Rad) were used for SDS-PAGE, with subsequent transfer onto a PVDF-membrane using a Trans-Blot SD Semi-Dry Transfer System (Bio-Rad). Subsequently, the membrane was incubated in 0.2% glutaraldehyde for 15 minutes, washed, and boiled in citrate antigen retrieval buffer (Vector) for 10 min in a 600W microwave oven. After cooling to room temperature, the membrane was incubated with a buffer containing 200 mM glycine for 10 min. Subsequent steps for Western blotting were identical to those listed above for the standard immunoblot protocol.

#### Quantitative RT-PCR

INS-1 cells (832/13) and mouse islets were washed with PBS, and total RNA was extracted using the RNeasy Mini Plus Kit for INS-1 cells and the RNeasy Micro Plus Kit for mouse islets (Qiagen), according to the manufacturer’s instructions. Total RNA was reverse-transcribed as described previously (Evans-Molina et al., 2007). Briefly, total RNA was incubated at 65°C for 5 min with 15 ng of random primers (Invitrogen) and 0.5 mM deoxynucleotide triphosphate (dNTP, Invitrogen). Next, 5x first-strand buffer (Invitrogen), 0.01 mM dithiothreitol (DTT, Invitrogen), and 200 U of Moloney murine leukemia virus reverse transcriptase (MMLV-RT, Invitrogen) were added to achieve a final reaction volume of 20 μL. The reverse transcription reaction was carried out at 37°C for 1 h and was followed by a denaturation step at 70°C for 15 min. Then, the resulting cDNA template was subjected to RT-qPCR using SensiFAST SYBR Lo-ROX reagents (Bioline) and a QuantStudio 3 thermocycler (Applied Biosystems). Relative RNA levels were established against β-actin mRNA using the comparative ΔΔCt method (Chakrabarti et al., 2002). Primer sequences are provided in the Key Resource Table.

#### Immunofluorescence and morphometric analysis

After euthanasia, the mouse pancreata were rapidly removed and fixed overnight using a buffered zinc formalin fixative Z-Fix (Anatech Ltd.). Fixed specimens were paraffin-embedded and longitudinal sections, 5 μm thick, were obtained. Following deparaffinization, rehydration, and antigen retrieval by heat in a citrate antigen retrieval buffer (Vector Laboratories), sections were permeabilized, blocked with Animal-Free Blocker (Vector Laboratories), and incubated with primary antibodies overnight at 4°C (Insulin – Dako; Cat# A0564; RRID:AB_10013624, Glucagon – Abcam; Cat# ab92517; RRID:AB_10561971). Incubation with secondary antibodies conjugated to AlexaFluor (Thermo Fisher) was performed at room temperature for 1 h. Sections were mounted with Fluorosave (Millipore). Images of islets were acquired using a Zeiss confocal microscope LSM 800 (Carl Zeiss, RRID:SCR_015963) using 40x and 63x oil immersion objectives. Images were analyzed with Zen software, blue edition (Carl Zeiss, RRID:SCR_013672).

For transmission electron microscopy (TEM), islets and INS-1 cells were fixed in 2% glutaraldehyde and 4% paraformaldehyde (Electron Microscopy Sciences) in 0.1 M sodium cacodylate buffer. TEM imaging was performed at the Advanced Electron Microscopy Facility at the University of Chicago (Chicago, IL).

The mass of β and α cells was estimated for each animal by determining the average β cell and α cell fractional area multiplied by the pancreatic weight as previously described (Jetton et al., 2005). In brief, five sections per mouse, each separated by ~25 μm, were immunostained for insulin and glucagon. Sections were deparaffinized, rehydrated, treated with hydrogen peroxide (Thermo Fisher Scientific) for 30 min to remove endogenous peroxidase activity, and subjected to antigen retrieval by heating in citrate buffer solution (Vector Laboratories). Subsequently, the slides were incubated with primary antibodies against insulin (Abcam, Cat# ab181547; RRID:AB_2716761) and glucagon (Abcam, Cat# ab92517; RRID:AB_10561971) overnight at 4°C, followed by incubation with horseradish peroxidase (HRP)-conjugated secondary antibody (NovaRed) at room temperature for 2 h. The color was developed using HRP substrate according to the manufacturer’s protocol (NovaRed), and the sections were counterstained with hematoxylin (Sigma-Aldrich). Slides were mounted with Permount mounting medium (Fisher) and scanned using AxioScan Z1 scanning microscope (Carl Zeiss, RRID:SCR_002677). The fractional area of β and α cells was measured using the ZEN 2 software (Carl Zeiss, RRID:SCR_013672).

#### Tandem mass tag MS/MS proteomic analysis of INS-1 cells

##### Sample Preparation

Four biological replicates of wild type and KO INS-1 cells (832/13) were lysed with 250 µL of 8M urea in 100 mM Tris HCl, pH 8.5. Following Bradford assay for protein quantitation (Protein Assay Dye Reagent Concentrate, Bio-Rad), 80 µg of each protein sample was reduced with 5 mM tris (2-carboxyethyl) phosphine hydrochloride (TCEP, Sigma-Aldrich) for 30 minutes at room temperature and alkylated with 10 mM chloroacetamide (CAA, Sigma Aldrich) for 30 min at room temperature in the dark. The reduced and alkylated protein samples were diluted in 50 mM Tris HCl, pH 8.5, to bring the concentration of urea below 2M. Samples were then digested overnight at 35ºC using Trypsin/Lys-C (Mass Spectrometry grade, Promega Corporation; enzyme-substrate ratio of 1:70). Digestion was quenched with trifluoracetic acid (TFA, 0.5% v/v, Fluka Analytical) and the products were desalted on Waters Sep-Pak cartridges (Waters) with a wash of 1 mL 0.1% TFA followed by elution in 50% and 70% acetonitrile (Fisher Chemical) containing 0.1% formic acid (Fluka Analytical) in the same collection tube. The elutes were dried and stored at −20ºC.

##### TMT labeling

Dried peptides were reconstituted in 29 µL of 50 mM triethylammonium bicarbonate, pH 8.0 (TEAB) (Sigma Life Science). A quantitative peptide assay (Quantitative Colorimetric Peptide Assay, Pierce) was done to ensure equivalent sample labeling, and 40 µg of peptides from each sample were labeled for 2 h at room temperature with 0.2 mg of Tandem Mass Tag (TMT) reagent (TMT™ Isobaric Label Reagent Set, Thermo Fisher Scientific). Four WT samples were labeled with TMT −126, −127N, −127C, −128N and four STIM1 KO samples were labelled with TMT −128C, −129N, 129C, and 130N respectively. Labeling reactions were quenched by adding 0.2% hydroxylamine (final v/v, Thermo Scientific) to the reaction mixture at room temperature for 15 minutes. Labeled peptides were then pooled and dried. The dried sample was reconstituted in 0.1% TFA and half was fractionated using a Waters Sep-Pak cartridge (Waters) with a wash of 1 mL 0.1% TFA and 0.5% acetonitrile (Fisher Chemical) containing 0.1% triethylamine (Thermo Scientific) followed by elution in 10%, 12.5%, 15%, 17.5%, 20%, 22.5%, 25%, and 70% acetonitrile (Fisher Chemical) containing 0.1% triethylamine.

##### Nano-LC-MS/MS Analysis

A tenth of each fraction was separated on a 25 cm Aurora column (AUR2-25075C18A, IonOpticks) at 400 nL/min in the EASY-nLC HPLC system (SCR: 014993, Thermo Fisher Scientific). The gradient used for the separation was 5-30% with mobile phase B for 160 min, 30-80% B over 10 min, and 80-10% B in the last 10 min (Mobile phases A: 0.1% FA, water; B: 0.1% FA, 80% Acetonitrile (Fisher Chemical). Nano-LC-MS/MS data were acquired in Orbitrap Eclipse™ Tribrid mass spectrometer (Thermo Fisher Scientific, RRID:SCR_022212) with a FAIMS pro interface. The mass spectrometer was operated in positive ion mode with advanced peak determination and Easy IC™ on and with 3 FAIMS CVs (−40, −55, −70); 1.3 sec cycle time per CV. Precursor scans (m/z 400-1650) were done with an orbitrap resolution of 120000, RF lens% 30, maximum inject time 105 ms, standard AGC target (4e5), MS2 intensity threshold of 2.5e4, including charges of 2 to 7 for fragmentation with 30 s dynamic exclusion. MS2 scans were performed with a quadrupole isolation window of 0.7 m/z, 38% HCD CE, 50000 resolution, 200% normalized AGC target (1e5), dynamic maximum IT, fixed first mass of 100 m/z. The data were recorded using the Xcalibur (4.3) software (Thermo Fisher Scientific, RRID:SCR_014593).

##### LC-MS Proteomics Data Analysis

RAW files were analyzed in Proteome Discover™ 2.5 (Thermo Fisher Scientific, RRID:SCR_014477) with a *Rattus norvegicus* UniProt reviewed and unreviewed FASTA and common contaminants (downloaded 2020_12_09, 35843 total sequences). SEQUEST HT searches were conducted with full trypsin, a maximum number of 3 missed cleavages, precursor mass tolerance of 10 ppm, and fragment mass tolerance of 0.02 Da. Static modifications used for the search were carbamidomethylation on cysteine (C) residues and TMT label on lysine (K) residues; Dynamic modifications used for the search were oxidation of methionines, TMT label on the N-termini of peptides, phosphorylation on serine/threonine/tyrosine, and deamidation of N (max 3 dynamic mods). Dynamic protein terminus modifications allowed were: acetylation (N-terminus), Met-loss, or Met-loss plus acetylation (N-terminus). Percolator False Discovery Rate was set to a strict setting of 0.01 and a relaxed setting of 0.05, and the Protein FDR validator in the consensus was set to a strict 1% protein FDR cutoff and relaxed 5% protein FDR cutoff. Co-isolation thresholds of 50% and average reporter ion S/N cutoffs of 5 were used for quantification. Lot-specific isotopic impurity correction levels were used and abundances were normalized to the total peptide amount. Normalized abundance values for each sample type, abundance ratio, log2(abundance ratio) values, and respective p-values (t-test) from Proteome Discover™ were exported to Microsoft Excel.

#### mRNA sequencing, library generation, and data analysis

Following 8 weeks of HFD treatment, islets were isolated from control (n=4) and STIM1Δβ (n=5) female mice, and bulk RNA sequencing analysis was performed. Briefly, after 24 h of recovery following islet isolation, total RNA was extracted from >100 hand-picked islets using the RNeasy Micro Kit (Qiagen). The quantity and quality of extracted RNA were evaluated using Agilent Bioanalyzer 2100 (Agilent, RRID:SCR_019389). The RNA integrity number (RIN) value of the samples was 9.378±0.342 (mean±SD). One hundred nanograms of total RNA was used to prepare a cDNA library. The preparation included mRNA purification/enrichment, RNA fragmentation, cDNA synthesis, ligation of index adaptors, and amplification. The protocols were performed according to the KAPA mRNA Hyper Prep Kit Technical Data Sheet, KR1352 – v4.17 (Roche). Each resulting indexed library was quantified, its quality was assessed by Qubit BioAnalyzer (Agilent, RRID:SCR_019715), and multiple libraries were pooled in equal molarity. The pooled libraries were then denatured, neutralized, and loaded to NovaSeq 6000 sequencer (Illumina, RRID:SCR_016387) at 300 pM final concentration for 100b paired-end sequencing. Approximately 30M reads per library were generated. A Phred quality score (Q score) was used to measure the quality of sequencing. More than 90% of the sequencing reads reached Q30 (99.9% base call accuracy).

The quality of sequencing data was first assessed using FastQC (Babraham Bioinformatics, RRID:SCR_014583). Data analysis was performed using Partek Flow Software Version 9.0.20.0202 (Partek, RRID:SCR_011860). Sequencing reads were aligned to mouse genome mm10 using STAR aligner version 2.6.1d. Transcripts with at least a total of 10 read counts were considered for downstream analysis. The aligned reads were annotated using RefSeq Transcripts 93 database. DESeq2 was used to identify differentially expressed RNAs (linear scale fold-change, FC ≥ 1.5 and p-value < 0.05). Log cpm values were used to generate PCA plots and heat maps.

Functional enrichment analysis, including pathway analysis and identification of upstream regulators, was performed using the Ingenuity Pathway Analysis tool (Qiagen, RRID:SCR_008653). Only pathways with a p-value < 0.05 were considered significant. We focused on the estrogen-related terms (beta-estradiol, estrogen, estrogen receptor, ESR1, ESR2, and ESRRA) for upstream regulators and identified the downstream genes predicted to be regulated by the aforementioned upstream regulators. Gene ontology functions (focusing on Biological process with p-value < 0.05) of the downstream genes were identified using Database for Annotation, Visualization, and Integrated Discovery (DAVID) (Huang da et al., 2009a, b).

#### Quantification of intracellular cAMP levels

Intracellular cAMP levels in INS-1 cells were measured using the cAMP - Gs Dynamic Homogeneous Time-Resolved Fluorescence (HTRF) kit (CisBio), following the manufacturer’s protocol. Briefly, 75,000 cells per well were seeded onto a 384-well white culture plate (Perkin Elmer). The next day, cells were incubated in an assay buffer consisting of HBSS supplemented with 20 mM HEPES (Gibco) and then lysed using cAMP-cryptate (Cisbio) and anti-cAMP-d2 antibody solutions (Cisbio). Lysed samples were incubated at room temperature for 1 h, and the plate was read on a SpectraMax iD5 multi-mode microplate reader (Molecular Devices) using excitation at 314 nm and emission at 620 and 668 nm. The cAMP concentration was determined through a standard curve interpolation using log(inhibitor) versus response – variable slope (four parameters) setting in Prism 9 (GraphPad, RRID:SCR_002798).

#### siRNA transfection

INS-1 (832/13) cells were transfected with siRNA targeted to *Gper1* using Lipofectamine 3000 (Invitrogen) according to the manufacturer’s recommendations. Approximately 0.5 × 10^6^ cells were plated in each well of a 12-well plate and immediately transfected with either 50 nM of control RNA (Ambion) or a mixture of three siRNAs targeting Gper1 gene (ID# s139733, s139734, s139735; Ambion) diluted in Lipofectamine 3000 solution (Invitrogen) and OptiMEM1 medium at 37°C. Cells were cultured for an additional 48 hours in the INS-1 cell culture medium, then harvested for total RNA isolation using the RNeasy Mini Plus Kit (Qiagen) and RT-qPCR analysis using the method outlined above.

#### Treatment with GPER1 agonist E_2_ and antagonist G-15

INS-1 (832/13) cells were plated in a 12-well plate at a density of approximately 0.5 × 10^6^ cells per well and incubated in 1 mL of the phenol red-free RPMI1640 medium (Gibco) at 37°C in the presence of 5% CO_2_. After 48 hours, the medium was removed and replaced with either fresh medium, or medium containing 0.1 μM of GPER1 agonist E_2_ (Steraloids) or 15 μM of GPER1 antagonist G-15 (Fan et al., 2018). Cells were then cultured for an additional 24 hours and collected for total RNA isolation using the RNeasy Mini Plus Kit (Qiagen) and RT-qPCR analysis using the method outlined above. The same protocol, employing 75 islets per well in a 6-well plate, was used for experiments with mouse islets.

### QUANTIFICATION AND STATISTICAL ANALYSIS

Statistical analysis was performed using the Prism 8.2.0 software (GraphPad, RRID:SCR_002798). To compare two data sets, the Student’s t-test was used. For experiments involving 3 independent groups, data were analyzed by a one-way ANOVA. Glucose and insulin tolerance tests were analyzed by a two-way ANOVA followed by the Tukey multiple comparison test. Results are reported as the mean ± SEM. A p-value of <0.05 was considered to indicate a significant difference between groups. Statistical analyses for proteomics and RNA sequencing are detailed above.

**KEY RESOURCES TABLE (see attached file)**

