## Supplementary material for "Stromal Interaction Molecule 1 Maintains β Cell Identity and Function in Female Mice through Preservation of G Protein-Coupled Estrogen Receptor 1 Signaling": Key Resources Table

The table highlights the genetically modified organisms and strains, cell lines, reagents, software, and source data **essential** to reproduce results presented in the manuscript. Depending on the nature of the study, this may include standard laboratory materials (i.e., food chow for metabolism studies), but the Table is **not** meant to be comprehensive list of all materials and resources used (e.g., essential chemicals such as SDS, sucrose, or standard culture media don’t need to be listed in the Table). **Items in the Table must also be reported in the Method Details section within the context of their use.** The number of **primers and RNA sequences** that may be listed in the Table is restricted to **no more than ten each**. If there are more than ten primers or RNA sequences to report, please provide this information as a supplementary document and reference this file (e.g., See Table S1 for XX) in the Key Resources Table.

***Please note that ALL references cited in the Key Resources Table must be included in the References list.*** Please report the information as follows:

- **REAGENT or RESOURCE:** Provide full descriptive name of the item so that it can be identified and linked with its description in the manuscript (e.g., provide version number for software, host source for antibody, strain name). In the Experimental Models section, please include all models used in the paper and describe each line/strain as: model organism: name used for strain/line in paper: genotype. (i.e., Mouse: OXTR^fl/fl^: B6.129(SJL)-Oxtr^tm1.1Wsy/J^). In the Biological Samples section, please list all samples obtained from commercial sources or biological repositories. Please note that software mentioned in the Methods Details or Data and Software Availability section needs to be also included in the table. See the sample Table at the end of this document for examples of how to report reagents.
- **SOURCE:** Report the company, manufacturer, or individual that provided the item or where the item can obtained (e.g., stock center or repository). For materials distributed by Addgene, please cite the article describing the plasmid and include “Addgene” as part of the identifier. If an item is from another lab, please include the name of the principal investigator and a citation if it has been previously published. If the material is being reported for the first time in the current paper, please indicate as “this paper.” For software, please provide the company name if it is commercially available or cite the paper in which it has been initially described.
- **IDENTIFIER:** Include catalog numbers (entered in the column as “Cat#” followed by the number, e.g., Cat#3879S). Where available, please include unique entities such as [RRIDs](https://www.force11.org/group/resource-identification-initiative), Model Organism Database numbers, accession numbers, and PDB or CAS IDs. For antibodies, if applicable and available, please also include the lot number or clone identity. For software or data resources, please include the URL where the resource can be downloaded. Please ensure accuracy of the identifiers, as they are essential for generation of hyperlinks to external sources when available. Please see the Elsevier [list of Data Repositories](https://www.elsevier.com/authors/author-resources/research-data/data-base-linking) with automated bidirectional linking for details. When listing more than one identifier for the same item, use semicolons to separate them (e.g. Cat#3879S; RRID: AB_2255011). If an identifier is not available, please enter “N/A” in the column.
  - ***A NOTE ABOUT RRIDs:*** We highly recommend using RRIDs as the identifier (in particular for antibodies and organisms, but also for software tools and databases). For more details on how to obtain or generate an RRID for existing or newly generated resources, please [visit the RII](https://www.force11.org/group/resource-identification-initiative) or [search for RRIDs](https://scicrunch.org/resources).

Please use the empty table that follows to organize the information in the sections defined by the subheading, skipping sections not relevant to your study. Please do not add subheadings. To add a row, place the cursor at the end of the row above where you would like to add the row, just outside the right border of the table. Then press the ENTER key to add the row. Please delete empty rows. Each entry must be on a separate row; do not list multiple items in a single table cell. Please see the sample table at the end of this document for examples of how reagents should be cited.

***TABLE FOR AUTHOR TO COMPLETE***

*Please upload the completed table as a separate document.* ***Please do not add subheadings to the Key Resources Table.*** *If you wish to make an entry that does not fall into one of the subheadings below, please contact your handling editor. (****NOTE:*** *For authors publishing in Current Biology, please note that references within the KRT should be in numbered style, rather than Harvard.)*

**KEY RESOURCES TABLE**

| REAGENT or RESOURCE | SOURCE | IDENTIFIER |
| --- | --- | --- |
| **Antibodies** | | |
| Mouse monoclonal anti-Actin (1:10000) | Millipore | Cat# MAB1501; RRID:AB_2223041 |
| Guinea pig polyclonal anti-Insulin (1:500) | Dako | Cat# A0564; RRID:AB_10013624 |
| Rabbit monoclonal anti-Insulin (1:500) | Abcam | Cat# ab181547; RRID:AB_2716761 |
| Rabbit monoclonal anti-Glucagon (1:500) | Abcam | Cat# ab92517; RRID:AB_10561971 |
| Mouse monoclonal anti-STIM1 (1:1000) | BD Biosciences | Cat# 610954; RRID:AB_398267 |
| Rabbit polyclonal anti-GPER1 (1:1000) | Abcam | Cat# ab39742; RRID:AB_1141090 |
| IIRDye® 800CW Donkey anti-Mouse IgG Secondary Antibody (1:10000) | LI-COR Bioscience | Cat# 926-32212; RRID:AB_621847 |
| IRDye® 680RD Donkey anti-Rabbit IgG Secondary Antibody (1:10000) | LI-COR Bioscience | Cat# 926-68073; RRID:AB_10954442 |
| Donkey anti-guinea pig IgG Secondary Antibody (Alexa Fluor-647) (1:10000) | Jackson ImmunoResearch Laboratories, Inc | **Cat# 706-605-148; RRID:** AB_234047 |
| Goat anti-rabbit IgG Secondary Antibody (Alexa Fluor-488) (1:10000) | Invitrogen | Cat# **A-11034; RRID:** AB_2576217 |
| Horseradish Peroxidase (HRP)-conjugated secondary antibody, anti-rabbit | NovaRed | Cat# PI-1000-1 |
| Horseradish Peroxidase (HRP) Substrate | NovaRed | Cat# SK-4800 |
| Anti-cAMP-d2 Antibody Solutions | Cisbio | Cat# 62AM4PEB |
| **Biological Samples** | | |
| βSTIM1KO mouse Islets and FFPE pancreas sections | This paper | N/A |
| C57BL6/J mouse Islets and FFPE pancreas sections | This paper | N/A |
| **Chemicals, Peptides, and Recombinant Proteins** | | |
| Insulin (regular) | Novo Nordisk | Cat# NC0769896 |
| D-Glucose | Sigma | Cat# G7528 |
| RPMI1640 without phenol red | Gibco | Cat#11835030 |
| Fetal bovine serum, charcoal-stripped | Millipore Sigma | Cat# F6765-500ML |
| 17β-Estradiol | Steraloids | Cat# E0950 |
| G-1 | Cayman | Cat# 1008933 |
| G-15 | Cayman | Cat# 14673 |
| ML-9 | Santa Cruz | Cat# sc-200519 |
| 2-APB | Tocris | Cat# 1224/10 |
| AncoA4 | Millipore | Cat# 532999 |
| Thapsigargin (TG) | Tocris | Cat# 1138/1 |
| Tunicamycin (TM) | Tocris | Cat# 3516/10 |
| Glucosamine | ChemCruz | Cat# sc-211202 |
| Random Primers | Invitrogen | Cat# 48190011 |
| Deoxynucleotide Triphosphate (dNTP) | Invitrogen | Cat# 10297018 |
| 5x First-Strand Buffer | Invitrogen | Cat# R1362 |
| Dithiothreitol (DTT) | Invitrogen | Cat# D1532 |
| Moloney Murine Leukemia Virus Reverse Transcriptase (MMLV-RT) | Invitrogen | Cat# 28025013 |
| SensiFAST SYBR Lo-ROX Reagents | Bioline | Cat# BIO-94050 |
| Chloroacetamide (CAA) | Sigma-Aldrich | Cat# C0267 |
| Trypsin/Lys-C (Mass Spectrometry grade, Promega Corporation; enzyme-substrate ratio of 1:70) | Promega Corporation | Cat# V5071 |
| Acetonitrile | Fisher Chemical | Cat# A955-4 |
| Formic Acid (FA) | Fluka Analytical | Cat# 56302-50ML-GL |
| Triethylammonium bicarbonate (TEAB) | Sigma Life Science | Cat# T7408-100ML |
| Tandem Mass Tag (TMT) Reagent (TMT Isobaric Label Reagent Set) | Thermo Fisher Scientific | Cat# A37725 |
| 50% Hydroxylamine | Thermo Scientific | Cat# 90115 |
| Triethylamine (0.1%) | Thermo Scientific | Cat# 1862986 |
| HBSS, no Ca no Mg no Phenol red | Gibco | Cat# 14175095 |
| HEPES | Gibco | Cat# 15630080 |
| cAMP-Cryptate | Cisbio | *Cat#* 62AM4PEB |
| Negative Control siRNA #1 | Ambion | Cat# AM4611 |
| Silencer Select siRNA (Gper1) | Ambion | Cat# 4390771 (ID# s139733, s139734, s139735) |
| Lipofectamine 3000 | Invitrogen | Cat# L3000001 |
| Paraformaldehyde | Electron Microscopy Sciences | Cat# 15710 |
| Antigen Unmasking Solution, Citrate-Based | Vector | Cat# H-3300-250 |
| **Critical Commercial Assays** | | |
| RNeasy Mini Kit | Qiagen | Cat# 74136 |
| RNeasy Micro Kit | Qiagen | Cat# 74034 |
| SsoAdvanced™ Universal SYBR® Green Supermix | Bio-Rad | Cat# 1725274 |
| *DC* Protein Assay | Bio-Rad | Cat# 5000112 |
| Insulin ELISA | Mercodia | Cat# 10-1247-10 |
| Glucagon ELISA | Mercodia | Cat# 10-1281-10 |
| Fura-2, AM, cell permeant | Invitrogen | Cat# F1221 |
| Calcium-6 | Molecular Devices | Cat# R8190 |
| cAMP Gs Dynamic Kit | Cisbio | Cat# 62AM4PEB |
| BCA Protein Quantification Assay | Bio-Rad | Cat# 5000001 |
| Bradford Assay for Protein Quantification | Protein Assay Dye Reagent Concentrate, Bio-Rad | Cat# 5000201 |
| Quantitative Colorimetric Peptide Assay | Pierce | Cat# 23275 |
| **Deposited Data** | | |
| RNA-seq raw data | This paper | GEO: Accession #GSE208135. |
| Mass spectrometry data | This paper | MassIVE with repository ID of MSV000089927. |
| **Experimental Models: Cell Lines** | | |
| STIM1 null INS-1 832/13 | Kono et al., Diabetes 2018 | N/A |
| INS-1 832/13 | Clark et al., Diabetes 1997 | RRID: CVCL_7226 |
| αTC1 Clone 9 | Hamaguchi et al., Diabetes 1990 | RRID: CVCL_0150 |
| **Experimental Models: Organisms/Strains** | | |
| Mouse: β cell specific STIM1-null (STIM1Δβ) in C57BL6/J background | This paper | N/A |
| Mouse: C57BL6/J | The Jackson Laboratory | Jax # 000664 |
| **Oligonucleotides** | | |
| Primer sequences |  |  |
| Mouse beta-actin-F | This paper | AGGTCATCACTATTGGCAACGA |
| Mouse beta-actin-R | This paper | CACTTCATGATGGATTGAATGTAGTT |
| Mouse slc2a2-F | This paper | GGCACAGACACCCCACTTAC |
| Mouse slc2a2-R | This paper | GCCAACATTGCTTTGATCCT |
| Mouse MafA-F | This paper | CCTGTAGAGGAAGCCGAGGAA |
| Mouse MafA-R | This paper | CCTCCCCCAGTCGAGTATAGC |
| Mouse MafB-F | This paper | TTCGACCTTCTCAAGTTCGACG |
| Mouse MafB-R | This paper | TCGAGATGGGTCTTCGGTTCA |
| Mouse UCN3-F | This paper | AGCACCCGGTACAGATACCAA |
| Mouse UCN3-R | This paper | GGCCTTGTCGATGTTGAAGAG |
| Mouse NKX6.1-F | This paper | CCTCTGGCCCGAACTCTGA |
| Mouse NKX6.1-R | This paper | GCTGCCACCGCTCGATT |
| Mouse PDX1-F | This paper | CGGCTGAGCAAGCTAAGGTT |
| Mouse PDX1-R | This paper | TGGAAGAAGCGCTCTCTTTGA |
| Mouse GPER-F | This paper | TCATTTCTGCCATGCACCCA |
| Mouse GPER-R | This paper | GTGGACAGGGTGTCTGATGT |
| Mouse/Rat Glucagon_F | This paper | TCACAGGGCACATTCACCAG |
| Mouse/Rat Glucagon_R | This paper | CATCATGACGTTTGGCAATGTT |
| Rat MafA-F | This paper | CCTGTAGAGGAAGCCGAGGAA |
| Rat MafA-R | This paper | CCTCCCCCAGTCGAGTATAGC |
| Rat MafB-F | This paper | ACCAAGGACGAGGTGATCC |
| Rat MafB-R | This paper | CAGGTGATGTTTCTGCTGGA |
| **Software and Algorithms** | | |
| Zen Blue edition ver2.3 | Carl Zeiss | https://www.zeiss.com/microscopy/int/products/microscope-software.html; RRID:SCR_013672 |
| Axio-Vision Software | Carl Zeiss | https://www.zeiss.com/microscopy/int/products/microscope-software.html; RRID:SCR_002677 |
| ImageJ | Fiji; Schneider et al., 2012 | Open source: <https://imagej.net/Fiji> RRID:SCR_002285 |
| Prism 8.2.0 | GraphPad Software | https://www.graphpad.com/;  RRID:SCR_002798 |
| Image Studio Software | LI-COR | RRID:SCR_015795 |
| Xcalibur (4.3) Software | Thermo Fisher Scientific | RRID:SCR_014593 |
| Proteome Discoverer 2.5 (*Rattus norvegicus* UniProt reviewed and unreviewed FASTA and common contaminants) | Thermo Fisher Scientific | RRID:SCR_014477 |
| FastQC | Babraham Bioinformatics | RRID:SCR_014583 |
| Partek Flow Software Version 9.0.20.0202 | Partek | RRID:SCR_011860 |
| STAR aligner version 2.6.1d | N/A | RRID:SCR_004463 |
| Ingenuity Pathway Analysis Tool | Qiagen | RRID:SCR_008653 |
| **Other** | | |
| Contour Glucometer | Bayer | Cat# 193715101 |
| Contour glucometer strips | Bayer | Cat# 193708050K7 |
| LSM 800 confocal imaging system | Carl Zeiss | RRID:SCR_015963 |
| Bioanalyzer 2100 | Agilent | Cat# G2939BA; RRID:SCR_019715 |
| Odyssey CLx scanner | LI-COR | RRID:SCR_014579 |
| EchoMRI-500 | EchoMRI | RRID:SCR_017104 |
| 60% kCal fat diet | Research Diets | Cat# D12492 |
| LabMaster Metabolism Research Platform | TSE Systems | N/A |
| SpectraMax iD5 Multi-Mode Microplate Reader | Molecular Devices | N/A |
| Mini-Protean Tetra Apparatus | Bio-Rad | Cat# 1658004EDU |
| Intercept Blocking Buffer | LI-COR | Cat# 927-70001 |
| Mini-PROTEAN Tris/Tricine Precast Gels, 16.5% | Bio-Rad | Cat# 4563063 |
| Trans-Blot SD Semi-Dry Transfer System | Bio-Rad | Cat# 1703940 |
| QuantStudio 3 Thermocycler | Applied Biosystems | Cat# A28567 |
| Buffered Zinc Formalin Fixative Z-Fix | Anatech Ltd. | Cat# 171 |
| NovaSeq 6000 Sequencer | Illumina | RRID:SCR_016387 |
| Waters Sep-Pak C18 1 cc Vac Cartridge | Waters | Cat# WAT054955 |
| Aurora Column (25 cm) | IonOpticks | Cat# AUR2-25075C18A |
| EASY-nLC HPLC System | Thermo Fisher Scientific | Cat# SCR:014993 |
| Orbitrap Eclipse Tribrid Mass Spectrometer (FAIMS pro interface) | Thermo Fisher Scientific | Cat# FSN04-10000; RRID:SCR_022212 |
| Agilent Bioanalyzer 2100 | Agilent | RRID:SCR_019389 |
| Qubit BioAnalyzer | Agilent | RRID:SCR_019715 |
| 384-well white culture plate | Perkin Elmer | Cat# 6007680 |
